# Hoplonemertean larvae are planktonic predators that capture and devour active animal prey

**DOI:** 10.1101/2021.02.02.429399

**Authors:** George von Dassow, Cecili Mendes, Kara Robbins, Sonia Andrade, Svetlana Maslakova

## Abstract

The superficially-simple ciliated planktonic larvae of hoplonemerteans have been assumed to be lecithotrophic direct developers, even though many develop from such small eggs that it is hard to imagine how they could give rise to a viable juvenile without some phase of larval feeding. Indeed, attempts to raise such larvae to settlement without food invariably fail. Observations that some hoplonemertean larvae are found in plankton samples at a range of sizes, and much larger than hatchlings, suggests they must indeed feed somehow. Since these “planuliform” larvae lack apparent means to concentrate suspended algae or other unicellular food, one alternative hypothesis is that they are planktonic predators that hunt large prey. Here we provide direct evidence that this is indeed the case for six distinct species of hoplonemerteans. We recorded wild-caught larvae of *Paranemertes californica, Paranemertes* sp., *Gurjanovella littoralis, Emplectonema viride, Carcinonemertes epialti*, and *Ototyphlonemertes sp.* attacking, subduing, and devouring pelagic crustaceans, including barnacle nauplii, cyprids, copepods and their nauplii, and others. While there is no doubt that some hoplonemerteans are genuine lecithotrophs, our evidence suggests that many species in this group both feed and grow during an extended planktonic larval period. This conclusion has important consequences for biogeographic and life-history studies in this group, because it implies enhanced potential for long-distance dispersal. More broadly, the possibility that many animal larvae are actually carnivores invites reconsideration of prevailing stereotypes about metazoan developmental modes and the trade-offs between them.

## 1. Introduction

Nemerteans are spiralians that live in nearly all marine habitats wherein they actively hunt animal prey. Nemertean adults are among the top predators in benthic marine communities, and most species have planktonic larvae. Modern taxonomy divides phylum Nemertea into three species-rich classes, the Palaeonemertea, the Hoplonemertea, the Pilidiophora (Thollesson and Norenburg, 2003; Andrade et al., 2012, 2014; Kvist et al., 2014; Strand et al., 2019). The Pilidiophora are named for the long-lived planktotrophic pilidium larva which develops on a diet of phytoflagellates; this larval form with its maximally-indirect developmental mode is not directly relatable to other invertebrate larval forms, but rather is a novel invention of the pilidiophoran nemerteans (Maslakova 2010a, b; von Dassow et al., 2013). Some paleaonemerteans, however, develop via a “hidden trochophore” that shares clear traits with spiralian trochophores (Maslakova et al., 2004a,b). Rather than giving rise to a trochophore-like planktotroph, these become so-called planuliform larvae – uniformly-ciliated, elongated swimmers, many of which grow in the plankton by feeding on large animal prey (Maslakova and Hiebert, 2014).

Hoplonemerteans also develop via planuliform planktonic larvae, but with no clear vestiges of trochophore-like development. Hoplonemerteans have been assumed to develop directly, albeit with a swimming dispersal phase, and indeed some lay moderately large (>200 microns), yolky eggs that give rise to nearly the full suite of adult anatomical characters without feeding (Norenburg and Stricker, 2002; Maslakova and von Döhren, 2009). But some hoplonemerteans also clearly lay small eggs, in the 100 micron range, seemingly too small to give rise to a viable predatory juvenile (e.g. Delsman, 1915; Stricker and Reed, 1981; Hiebert et al., 2010). Also, phylogeographic studies indicate levels of population connectivity compatible with species possessing long-lived larvae (e.g. Tulchinsky et al. 2012; Andrade et al., 2011; Mendes et al., 2018). Furthermore, hoplonemertean “larvae” are routinely found in plankton samples, at sizes (and spectrum of sizes) that imply some kind of extensive feeding and growth (Maslakova and Hiebert, 2014).

This is especially stark in the case of the egg parasite *Carcinonemertes epialti*, which as an adult lives on crabs, feeding on their brooded eggs, amongst which it also lays cocoons of ~80 micron eggs. These eggs hatch as uniformly-ciliated, actively-swimming bullet-shaped “planuliform” larvae. Attempts to raise and settle these larvae onto their host species without food invariably failed (Roe, 1979; Stricker and Reed, 1981; Dunn and Young, 2014). Yet these are recovered in plankton samples (and unambiguously identified by DNA barcoding) at sizes up to 1 mm, implying greater than 1000-fold increase in mass (Maslakova and Hiebert 2014, as *C. errans*). These undergo a distinct, though subtle, metamorphosis upon settlement (Dunn and Young, 2014), and so qualify as larvae on morphological grounds. Many other hoplonemerteans, including species of *Emplectonema, Paranemertes, Ototyphlonemertes*, and *Poseidonemertes* are also recovered from plankton samples in a spectrum of sizes that imply feeding.

How hoplonemertean larvae might grow in the plankton has been a mystery. They lack apparent food-concentrating apparatus (e.g., ciliated bands or mucus houses) and do not accumulate algal food in culture. Occasional traces in DNA barcoding studies suggest that they may hunt large animal prey (Maslakova and Hiebert, 2014; and present study). Here we report that wild-caught hoplonemertean planktonic larvae actively capture and devour pelagic crustacean larvae and copepods. Furthermore, we report successful identification (by direct observation) of a prey acceptable to hatchling *Carcinonemertes*, and demonstrate growth thereon in the laboratory. Therefore, we conclude that the planuliform larvae of hoplonemerteans, despite their anatomical similarity to adults and similarity of feeding mechanism, occupy a distinct ecological niche from their parents, exploiting planktonic resources for growth in a manner – macrophagous carnivory – not widely recognized as a common strategy among invertebrate larvae.

## 2. Methods

### 2.1 Collecting hoplonemertean larvae

Larval nemerteans were recovered from regular dockside plankton tows in the Charleston Marina, Charleston, OR (43.344738° N 124.320947° W) using either a 150-or 53-micron mesh net, usually with a jar cod end, during Winter and Spring of 2012, 2019 and 2020. Most samples were collected near the surface shortly before or after high slack by towing the net back and forth between two adjacent slips. This seems to maximize recovery of nemertean larvae. Concentrated plankton was diluted with about the same volume of filtered natural seawater, and hand-sorted to retrieve individual larvae into bowls of clean filtered seawater. Samples were kept in seatables at 12–15 °C. Candidate prey, including various polychaetes, molluscs, and crustaceans were likewise retrieved from the same samples and rationed out to captive hoplonemertean larvae.

Hatchlings of *Carcinonemertes epialti* were obtained from crabs in berry (*Cancer productus, Cancer antennarius* and *Metacarcinus magister*) collected from nearby rocky shores. *Carcinonemertes’* cocoons were pulled with forceps from amongst the crab’s eggs, then kept in clean natural seawater in bowls at 12–15 °C, wherein they hatched and began to swim actively. Without food, these hatchlings swim for weeks, slowly dwindling in size.

### 2.2 Observations of feeding

Observations of feeding were initially made by using a stereomicroscope to follow individual larvae as they swam around their pens, observing them as they interacted with fellow captives. Once acceptable prey had been identified, video recordings were made by placing one or a few nemerteans and several freshly-collected prey items into a Syracuse dish with clean seawater, placed onto a darkfield stage beneath a Leica Z6 Apo macroscope. Recordings were made with a Point Grey Grasshopper 3 USB camera operated by StreamPix 7 (Norpix), which allows one to record a stream of frames into a circular buffer. Some feeding events were also documented on slide-and-coverslip cuvettes using an Olympus BX-51 DIC microscope and a Spot Insight CMOS camera.

### 2.3 Collecting crabs infested with *Carcinonemertes*

One gravid female of *Metacarcinus magister* was collected near Charleston, OR in November 2019 and kept in a tank with aeration until the eggs were harvested. A single individual of *Carcinonemertes* from this egg mass (COIMB-1929) was barcoded, and the sequence used in phylogenetic analysis (Supplemental Figure 1, also see Table 2 for Genbank accession number). Eggs of two other female crabs, one of *Cancer productus* and another of *Cancer antennarius* were obtained from the Charleston Marine Life Center (also collected near Charleston, OR) in February and April 2020. In all cases, crab eggs were removed with forceps and kept in 150 ml glass bowls with filtered seawater. These eggs were searched for *C. epialti* egg strands and adults. Once these were found, they were transferred to separate bowls with filtered seawater and checked daily for hatchlings. However, none of these *Carcinonemertes* individuals were barcoded. *Carcinonemertes* juveniles (but no reproductive adults, or egg sheathes) were also found on *Pugettia producta*, collected by SAM and CM intertidally at Sunset Bay, near Charleston, OR on October 15, 2019. One juvenile was barcoded and sequences used in the phylogenetic analysis (Supplemental Figure 1) and submitted to GenBank (Accession numbers: 16S MW520855, COI MW522615). Another *Carcinonemertes* individual was removed on August 17, 2016 from the invasive European green crab, *Carcinus maenas*, collected intertidally in Coos Bay, OR. This individual was also barcoded and sequences used in phylogenetic analysis (Supplemental Figure 1) and submitted to GenBank (Accession numbers: 16S MW520854, COI MW522616).

### 2.4 DNA barcoding of hoplonemertean larvae and their prey

Wild-caught hoplonemertean larvae were mounted between a glass slide and a coverslip supported by clay feet, and photographed using an Olympus BX-51 DIC microscope and a Leica DFC400 or a Spot Insight CMOS camera. After being photographed, larvae were retrieved from the slide and rinsed in microfiltered seawater, and frozen individually in a few microliters of sea water at −80°C. Individuals were either frozen immediately (and within ~30 min. of collection when attempting to amplify prey DNA from gut contents) or kept in bowls with filtered sea water and candidate prey, and frozen later after a specific observed feeding event in the lab. If anything remained of the prey, it was frozen and barcoded separately. DNA was extracted from individual larvae (or prey) using Instagene matrix (Biorad) following manufacturer’s protocol. Regions of Cytochrome Oxidase I or 16S rRNA were PCR-amplified using 500 nM each of a forward and reverse primer (Table 1), 1U of Go Taq DNA polymerase with provided buffer (Promega), 200 μM of dNTPs (NE Biolabs), and 2 μl of DNA extract, with the following parameters: initial denaturation at 95° C for 2 minutes, followed by 34 cycles of denaturation at 95° C for 40 seconds, primer annealing at 42-50° C for 40 seconds, extension at 72° C for 1 minute, and final extension at 72° C for 2 minutes. PCR products were evaluated using gel electrophoresis. Those containing single bright bands were purified using Wizard SV Gel and PCR Clean-up Kit (Promega), and sequenced in both directions using PCR primers at Sequetech Inc. (Mountain View, CA). Sequences were trimmed to remove primers and low quality ends, and assembled into contigs and compared against GenBank database using the BLASTn algorithm available on the NCBI website. COI sequences were checked for stop codons using Invertebrate Mitochondrial translation table. Sequence ID above 98% for 16S and above 95% for COI was considered to be a species-level match. Where no close matches were found in GenBank to permit species-level ID, sequences were identified to genus based on maximum likelihood trees (Supplemental Figures 1 and 2). Sequences are deposited in GenBank (see Table 2 for Accession numbers). The three top BLAST matches to each novel sequence were downloaded from GenBank and aligned with the novel sequences using the online version of Mafft v.7 (Katoh et al. 2019). The resulting alignments were input for Maximum Likelihood searches performed in RAxML v.8.2 (Stamatakis 2014) as available in the CIPRES Science Gateway online platform (Miller et al. 2010), with 1000 Bootstrap replicates and GTRGAMMA model.

**Table 1.**
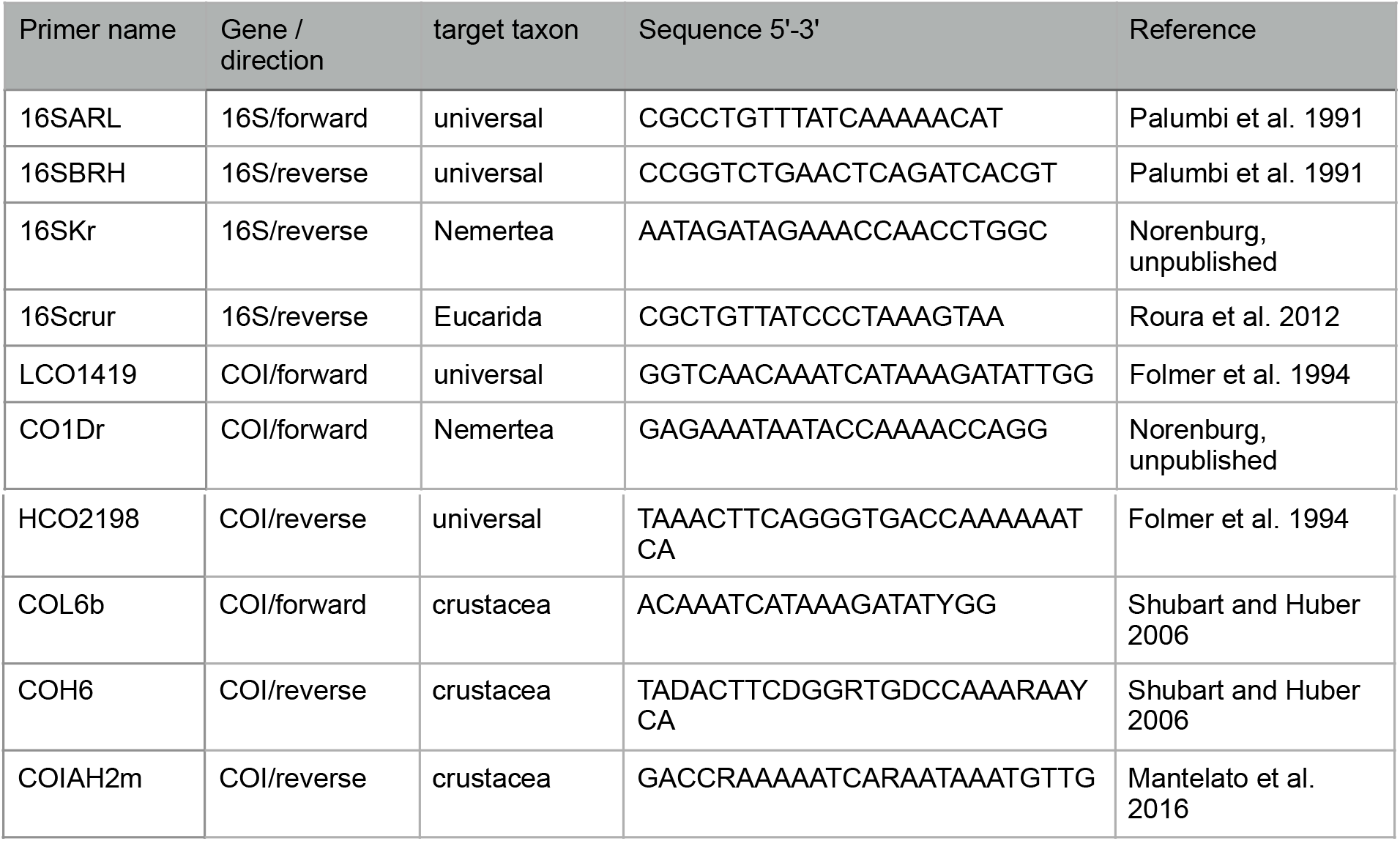
Primers used for DNA-barcoding of hoplonemertean larvae and their prey.

**Table 2.** Documented instances of predation by different species of hoplonemertean larvae on crustacean prey. Prey ID in bold represents instances of feeding in the wild prior to capture, detected by PCR, otherwise results are for predation observed in captivity. Prey without DNA barcodes was identified morphologically.

| Specimen identifier | Nemertean (predator) ID | Collecting information | Nemertean predator COI accession number | Crustacean prey ID | Crustacean prey COI accession number | Crustacean prey 16S accession number | Corresponding figures and videos of predation |
| --- | --- | --- | --- | --- | --- | --- | --- |
| N17 | *Paranemertes* sp. (*Paranemertes californica* species complex) | T. C. Hiebert, 5 Feb 2013, Charleston, OR, USA. | MW396772 | ***Scleroplax (Pinnixa) faba*** | - | MW403829 | - |
| N60 | *Paranemertes* sp. (*Paranemertes californica* species complex) | T.C. Hiebert, 19 Feb 2013, Charleston, OR, USA. | MW396771 | ***Cancer* sp.** | - | MW403832 | - |
| N97 | *Paranemertes* sp. (*Paranemertes californica* species complex) | T.C. Hiebert, 14 Mar 2013, Charleston, OR, USA. | MW396770 | ***Scleroplax (Pinnixa) faba*** | - | MW403830 | - |
| COIMB2046 Elvira | *Paranemertes* sp. (*Paranemertes californica* species complex) | C. Mendes and G. von Dassow, 2019, Charleston, OR, USA. | MW396769 | calanoid copepod nauplii and adults | - | - | - |
| 21III19-2 | *Paranemertes californica* | S.A. Maslakova, G. von Dassow, Mar 2019, Charleston, OR, USA. | MW396766 | *Centropages abdominalis* | MW401789 | - | Fig. 5A; Supplemental Video 5 |
| 21III19-5 | *Paranemertes californica* | S.A. Maslakova, G. von Dassow, Mar 2019, Charleston, OR, USA. | MW396761 | ***Scleroplax (Pinnixa) littoralis*** | - | MW403828 | - |
| 21III19-8 | *Paranemertes californica* | S.A. Maslakova, G. von Dassow, Mar 2019, Charleston, OR, USA. | MW396768 | *Carcinus maenas*; also calanoid copepod | - | MW403831 | Fig. 5B; Supplemental Video 6 |
| 8III19-1 | *Paranemertes californica* | S.A. Maslakova, G. von Dassow, Mar 2019, Charleston, OR, USA. | MW396765 | barnacle nauplius | - | - | - |
| COIMB204<br>9Victor | *Paranemertes californica* | C. Mendes and G. von Dassow, 2019, Charleston, OR, USA. | MW396767 | calanoid copepod | - | - | Figs. 1B, 2B; Supplemental Video 1 |
| COIMB211<br>7Hester | *Paranemertes californica* | C. Mendes and G. von Dassow, 2019, Charleston, OR, USA. | MW396764 | barnacle nauplius | - | - | Figs. 2A, A', 3B |
| COIMB204<br>8Anton | *Paranemertes californica* | C. Mendes and G. von Dassow, 2019, Charleston, OR, USA. | MW396763 | copepod nauplius | - | - | Fig. 3A |
| COIMB204<br>7Miriam | *Paranemertes californica* | C. Mendes and G. von Dassow, 2019, Charleston, OR, USA. | MW396762 |  | - | - |  |
| COIMB211<br>6Alfonse | *Gurjanovella littoralis* | C. Mendes and G. von Dassow, 2019, Charleston, OR, USA. | MW396773 | calanoid copepod nauplii and adults; barnacle nauplii; decapod zoeas; cladoceran | - | - | Figs. 2C,D, 4; Supplemental Videos 2–4 |
| 21III19-3 | *Emplectonema viride* | S.A. Maslakova, G. von Dassow, Mar 2019, Charleston, OR, USA. | MW396777 | barnacle nauplius | - | - | Fig. 6 |
|  | *Emplectonema viride* | C. Mendes, Feb 2020, plankton, Charleston, OR, USA | - | *Balanus glandula* | MW401788 | - | - |
|  | *Emplectonema viride* | C. Mendes, Feb 2020, plankton, Charleston, OR, USA | - | *Balanus glandula* | - | - | - |
|  | *Carcinonemertes epialti* | G. von Dassow, Jan 2020, plankton, Charleston, OR, USA | - | cyclopoid nauplius, copepodite | - | - | - |
| COIMB1929 | *Carcinonemertes epialti* | C. Mendes coll. January 2020 from egg mass of *Metacarcinus magister* coll. Nov 2019, Coos Bay, OR, USA. | MW396778 | *Oithona* sp. | MW401781 | - |  |
|  | *Carcinonemertes epialti* | C. Mendes coll. Feb 2020 from egg mass of *Cancer antennarius* coll. Feb 2020, Coos Bay, OR, USA | - | *Oithona* sp. |  |  | Fig. 8C |
|  | *Carcinonemertes epialti* | C. Mendes coll. January 2020 from egg mass of *Metacarcinus magister* coll. Nov 2019, Coos Bay, OR, USA. | - | *Oithona* sp. | MW401782 | - | - |
|  | *Carcinonemertes epialti* | C. Mendes coll. January 2020 from egg mass of *Metacarcinus magister* coll. Nov 2019, Coos Bay, OR, USA. | - | *Oithona* sp. | MW401783 | - | - |
|  | *Carcinonemertes epialti* | C. Mendes coll. January 2020 from egg mass of *Metacarcinus magister* coll. Nov 2019, Coos Bay, OR, USA. | - | *Oithona* sp. | MW401784 | - | - |
|  | *Carcinonemertes epialti* | C. Mendes coll. January 2020 from egg mass of *Metacarcinus magister* coll. Nov 2019, Coos Bay, OR, USA. | - | *Oithona* sp. | MW401785 | - | - |
|  | *Carcinonemertes epialti* | C. Mendes coll. January 2020 from egg mass of *Metacarcinus magister* coll. Nov 2019, Coos Bay, OR, USA. | - | *Oithona* sp. | MW401786 | - | - |
|  | *Carcinonemertes epialti* | C. Mendes coll. January 2020 from egg mass of *Metacarcinus magister* coll. Nov 2019, Coos Bay, OR, USA. | - | *Oithona* sp. | MW401787 | - |  |
| COIMB2113 | *Otothyphlonemertes* sp. | G. von Dassow, 6 June 2020, Charleston, OR, USA | MW396776 | copepod nauplius | - | - | Fig. 10 |
| COIMB2114 | *Otothyphlonemertes* sp. | G. von Dassow, 6 June 2020, Charleston, OR, USA | MW396774 |  |  |  |  |
| COIMB2115 | *Otothyphlonemertes* sp. | G. von Dassow, 6 June 2020, Charleston, OR, USA | MW396775 |  |  |  |  |

## 3. Results

### 3.1 Wild-caught planktonic hoplonemertean larvae catch and eat large animal prey

In contrast to pelagic grazers, animals which feed on relatively large prey would be expected to feed only rarely. Indeed, to observe feeding by hoplonemertean larvae it was necessary to capture several individuals, guess correctly their possible prey and supply it to them in excess, and then watch them continuously for several hours. Our first direct observation of feeding by a hoplonemertean larva was of a small, lentoid brownish-green specimen with four eyes, who was noticed nosing around a twitching but non-swimming barnacle nauplius of greater length. Over the course of ~10 min, the nemertean first broke into the cuticle, then sucked all soft tissue from within, leaving a nearly empty naupliar exoskeleton behind. The much-expanded nemertean departed from the husk of its prey, swimming actively for a time before coming to rest on the dish bottom.

Repeated observations of this individual and similar wild-caught hoplonemertean larvae confirmed unambiguously that these target crustacean prey, often much larger than themselves. Initial DNA barcoding attempts identified several of the wild-caught individuals as *Paranemertes californica* and *Emplectonema viride.* While *E. viride* larvae are morphologically distinctive and can be unambiguously identified in plankton samples, the morphotype that includes *P. californica* larvae, as will be detailed below, conflates at least three species that cannot readily be distinguished based on overt shape, color, stylet, or behavior. We grouped these in collections as “*Paranemertes-type*” larvae. Both *Emplectonema* and *Paranemertes-type* larvae subdue their prey with one or more strikes of the proboscis, then push their anterior end into a crack or hole in the cuticle, and suck out most or all of the soft tissue within (Fig. 1A). In many but not all instances the intrusion site matched one of the sites at which the proboscis (and presumably stylet) made contact (Fig. 1B; Supplemental Video 1). Following attacks, nemerteans are associated tightly with their catch and are not easily dislodged by water motion or even pipetting. However, transfer and cover-slipping usually induced nemerteans to abandon prey. There were some signs that mucus entanglement accompanied capture. However, it seemed clear that most of the *Paranemertes-type* (Fig. 2) and *Emplectonema viride* larvae envenomate their prey with some paralytic toxin. Small nauplii often ceased swimming immediately; copepods (which were typically much larger than the nemerteans) ceased escape swimming, though they often continued to twitch; zoeas (likewise usually larger than the nemerteans) ceased most movement except twitching, and the highly-visible heartbeat continued. Some nemerteans were successfully transferred to a slide-and-coverslip prep after capture, and a few actually captured prey within such a prep (Fig. 3). These cases allowed us to observe 1) that the nemertean entangles prey appendages with mucus (Fig. 3A, see inset frame 3), and 2) that prey tissue is at least partially disrupted (Fig. 3A,B), hypothetically by some proteolytic component of the venom injected during the proboscis strike, before the nemertean commences to drink the partially liquified tissues of its victim.

**Figure 1.**
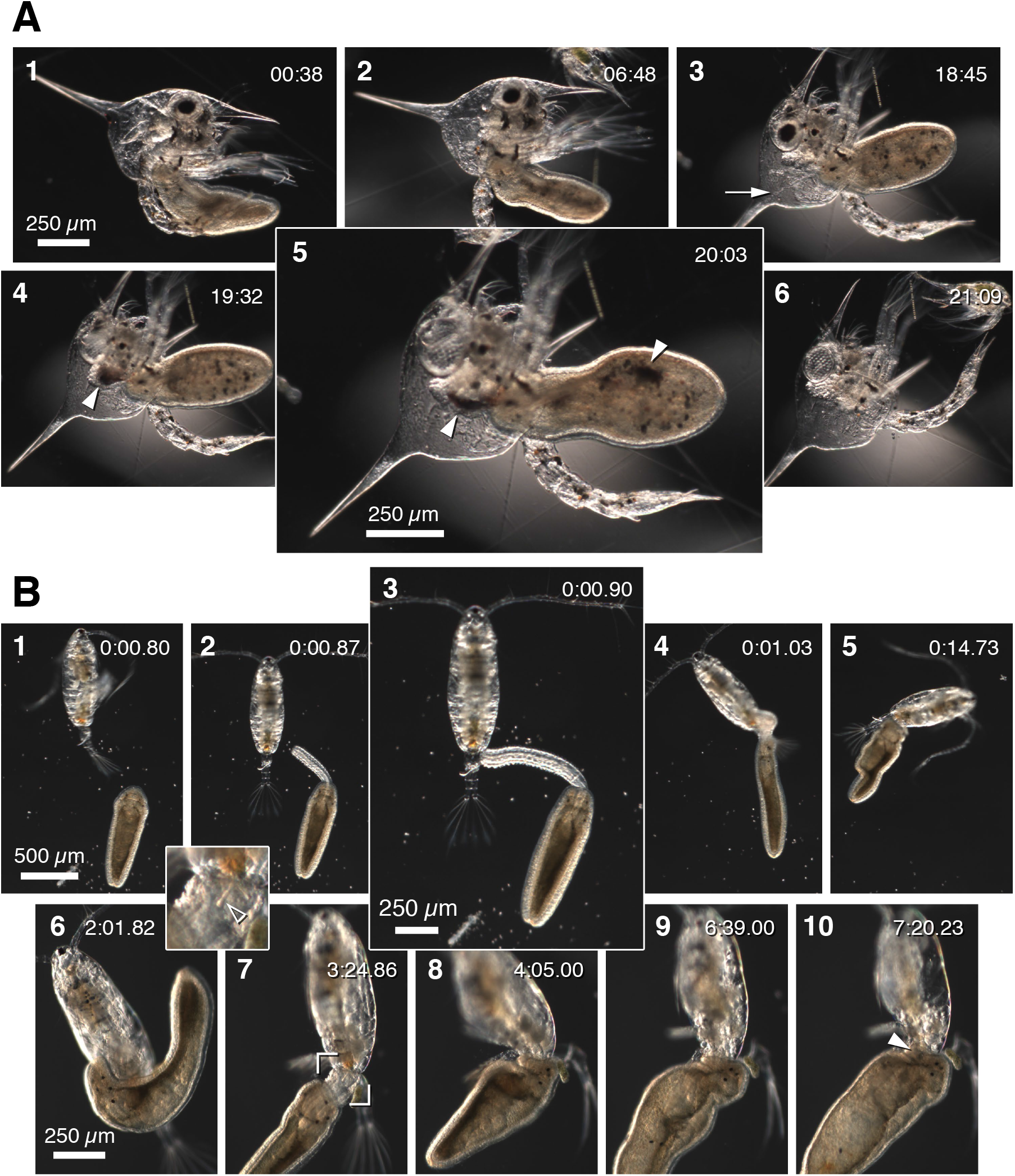
Pelagic hoplonemertean larvae catch and eat large, motile crustaceans. (A) Frames from an image sequence in which a wild-caught hoplonemertean larva <1 mm long devours a wild-caught decapod zoea. Recording began shortly after the initial attack; time is in min:sec from start of recording. After the initial attack, the nemertean apparently probed for an entry point for several minutes. During this time the zoea was alive but subdued. Within 15 min. the nemertean had broken into the cuticle posterior to the thorax and begun to ingest a stream of partially liquified tissue (arrow, frame 3, points out strand of tissue drawn from the dorsal spine). Arrowheads in frames 4 and 5 indicate the pigment of each zoeal eye; frame 5 is a 1.5x blowup relative to others. After ingesting most of the soft tissues of the head and thorax, the nemertean abandoned the carcass without consuming the tissues within the abdomen or appendages. (B) Frames from a video recording (corresponds to Supplemental Video 1), one of the few which documents a feeding event from start to finish and in which the nemertean was swimming at the time it captured its prey, in this case a calanoid copepod ~1 mm long. The top row shows the initial capture at one magnification; magnification increased in bottom row. Frame 3 is a 1.5x blowup compared to others, and shows the moment the proboscis struck the prey. Note from frames 2 and 3 that the nemertean must be able to aim its proboscis relative to its body axis. The nemertean remained attached to the prey, which attempted to escape, as it retracted its proboscis (frames 4,5). Subsequently, the worm apparently used its proboscis and stylet (hollow arrowhead; inset, frame 7, is a 3x blow-up) in successive strikes to penetrate the cuticle, whereafter it devoured most of the soft tissue. Time is in min:sec relative to start of recording. DNA barcoding showed that these two nemertean larvae belong to *P. californica* (see Table 2).

**Figure 2.**
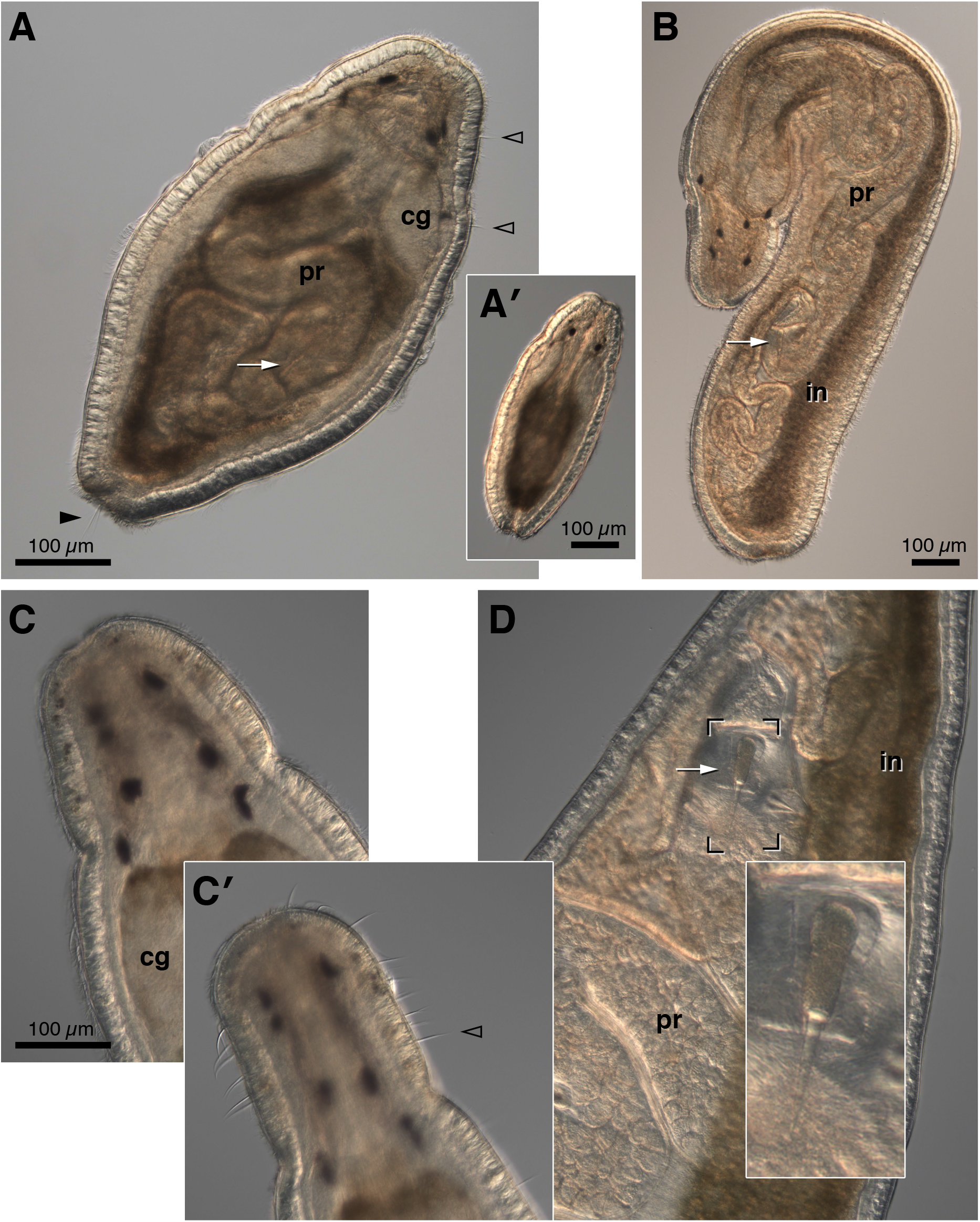
Wild-caught Paranemertes-type larvae, *Paranemertes californica* and *Gurjanovella littoralis*. Three individuals are shown, assigned nomens for sample-tracking purposes: “Hester” (A and A’), “Victor” (B) (both *P. californica*), and “Alfonse” (*G. littoralis*) (C, D). All were barcoded, as summarized in Table 1, and appear in other figures and supplemental videos included here. All were cultivated in the lab in isolated bowls for several weeks, during which time they were offered various prey, and underwent substantial growth. In the images shown here, Hester (A, A’) has already eaten a meal in the lab (Fig. 3B), but is nevertheless typical of size and stage at collection: ~500 μm and possessed of complete proboscis (pr) with stylet (arrow in A), 4-6 eyespots, and a few short ciliary cirri (hollow arrowheads) anterior to the cerebral ganglia (cg) in addition to a posterior cirrus (solid arrowhead). The larva in (A) is compressed by the coverslip to show proboscis and stylet; image of uncompressed larva (A’) emphasizes typical resting posture, including prominent mid-ventral groove. Victor (B) grew to ~2 mm long before preservation for DNA barcoding; as shown here this individual (who is featured in Fig. 1B and Supplemental Video 1) typifies large but still actively-swimming worms (in=intestine). Alfonse (C,D) was caught at relatively large size, and is depicted after several weeks in culture and many meals’ growth (see Fig. 4 and Supplemental Video 2–4), and at the point depicted no longer swam actively, but rather drifted on a thread or swam slowly along the bottom of a dish. Notably, these large post-swimming individuals have many ciliary cirri in the head region, which can be flattened or extended (C vs. C’); sparser cirri are also found along the body margin. At this stage these animals have an enormous papillate proboscis with ~50 μm main stylet and several accessory ones in adjacent pouches, >6 eyespots (some doubled), and often trail a visible thread from the very posterior. Inset in D is a 2.5x blowup of the stylet and basis.

**Figure 3.**
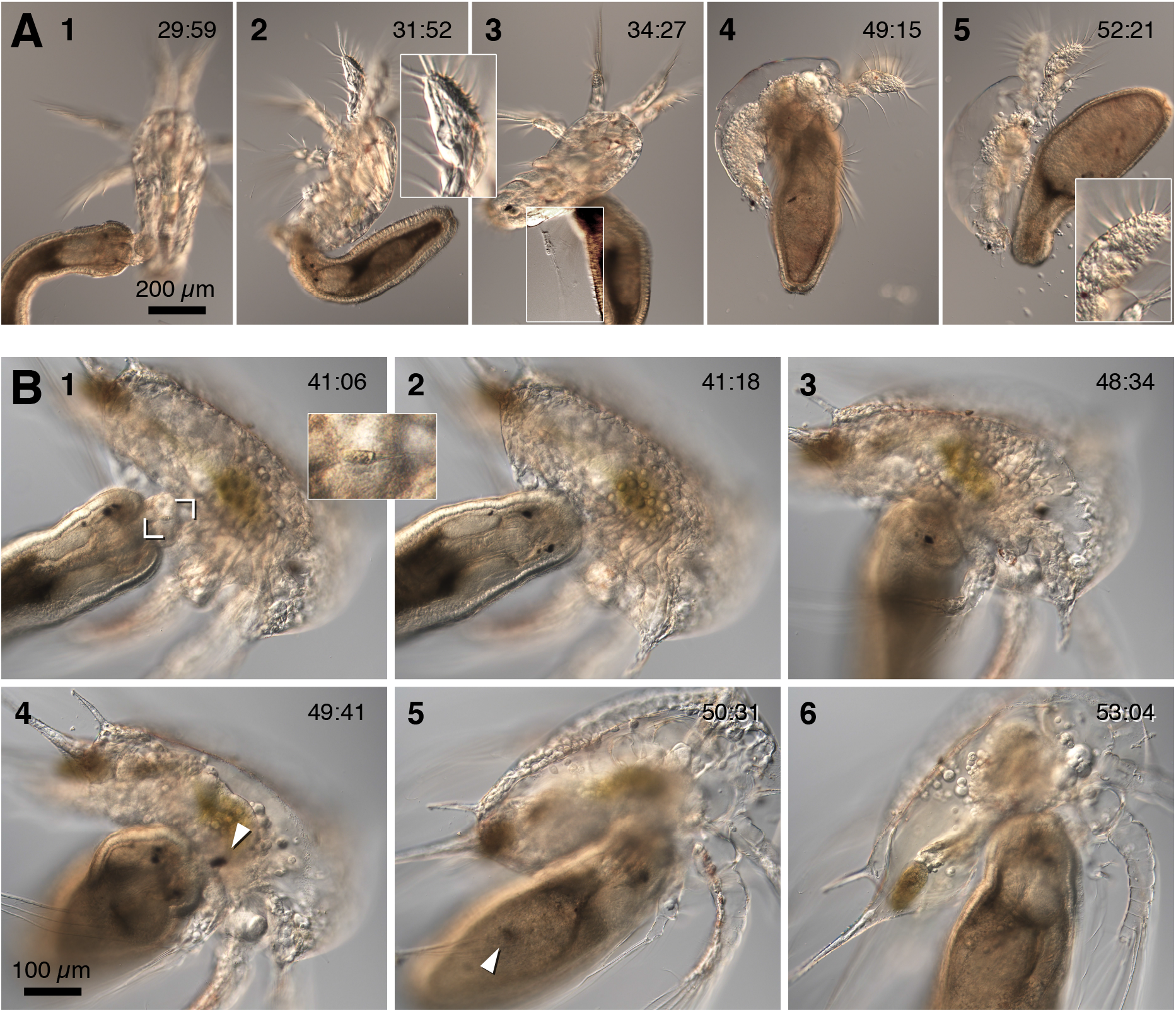
*Paranemertes californica* larvae entrap their prey in mucus and disrupt their tissues before consuming them. (A) Image sequence in which a small wild-caught individual (“Anton” in Table 1) attacks and devours a copepod nauplius. This sequence was recorded on a slide-and-coverslip prep with ample room; animals re-orient from frame to frame because they were free to move. Timestamps are clock time in min:sec. Frame 1 shows the end of the second strike, closely following the initial one in the same zone. Shortly thereafter, as the nemertean appeared to seek an entrance, the copepod’s tissue was clearly intact (inset is a 2.5x blowup of distal antennule segment; note also the strongly-birefringent dorsal muscles). In frame 3 the nemertean has shifted position as it explores the prey; in so doing, it becomes apparent that it has deposited a substantial clot of mucus near the site of the original strike (contrast-enhanced box in frame 3). Within 20 min., as the nemertean ingested the copepod tissue through a newly-made entrance hole in the ventral thorax, it became apparent that the prey tissue was almost totally disrupted. As the replete worm abandons the remains of its victim, comparison of the boxed inset (2.5x blowup of the distal antennule segment) emphasizes the extent of tissue disruption. (B) Image sequence in which a different individual (“Hester” in Table 1; Fig. 2A) devours a barnacle nauplius. As in (A), this sequence was recorded on a slide-and-coverslip prep with ample room for movement, and timestamps are clock time in min:sec. Frame 1 illustrates the most common site of penetration: just posterior and distal to the base of the mandible. Inset is a 3x blowup showing the stylet; this sequence begins with a secondary strike, shortly before the worm successfully penetrated the cuticle. Although the nauplius was initially still partly mobile and tissue appeared intact, once the worm began to ingest tissue (frame 3) the flesh of the nauplius was partially disrupted. Arrowheads in frames 4 and 5 indicate the naupliar eye before and after ingestion. Note how the nemertean empties even appendages of soft tissue (frames 4-6).

### 3.2 Capture strategy and feeding behavior

Observed capture and feeding events in large custard bowls took a variety of forms, the important categories being a) instances in which the more or less sedentary nemertean attacked an active, passing prey item with a proboscis strike (Fig. 4 and Supplemental Video 2– 4), b) instances in which the errant nemertean came upon a trapped or motionless prey item near the glass or water surface and attacked it after a brief investigation (Fig. 3), and c) cases in which the errant and actively-swimming nemertean attacked actively-swimming prey (Fig. 1B and Supplemental Video 1). For obvious reasons the latter category is difficult to record. Most recordings of capture and feeding took place in small, brightly-lit shallow glass arenas (Syracuse dishes) in which it seems clear the nemerteans are not particularly comfortable, and most successful recordings were of the first category: nemerteans had ceased active swimming, crawling slowly on the bottom; an active crustacean approached within striking distance, and its motion apparently triggered a strike.

**Figure 4.**
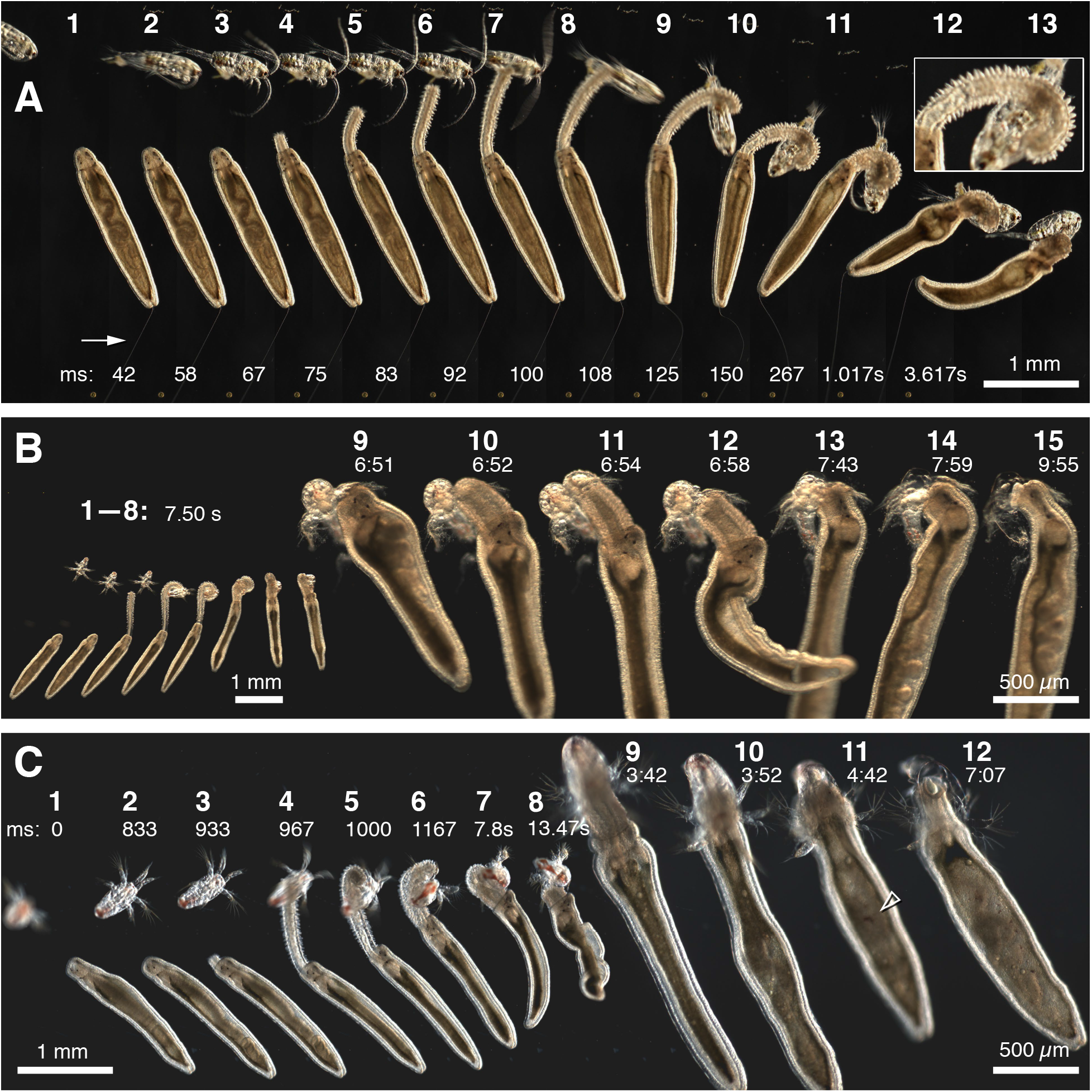
Prey strikes by *Gurjanovella litorallis*. All sequences shown in this figure feature Alfonse (Fig. 2C,D; Table 1) and correspond to Supplemental Videos 2-4. (A) Montage of frames from a high-speed (120 fps) recording in which the nemertean caught a passing calanoid copepod (corresponds to Supplemental Video 2). Note the visible thread (arrow) trailing from the posterior end of the worm. Times in ms are roughly registered with this thread. At the start of the sequence the copepod swims to within 1 mm of the worm’s anterior end, and pauses. The proboscis strike took less than 30 ms to contact the copepod (frames 4-7); note that the bend in the proboscis as it everts implies that the nemertean can aim it (as in Fig. 1B and Supplemental Video 1). After being struck, the copepod promptly attempted to jump (frame 8: note motion blur and copepod’s folded antennae and extended thoracopods). After a few seconds (frames 12,13) the nemertean retracted its proboscis, remaining attached to the prey, presumably by entangling mucus. Note the prominent papillae on the everted proboscis: inset is a 2x blowup taken from frame 10. (B) Another sequence involving a calanoid nauplius. Timestamps here are in min:sec relative to start of recording (corresponds to Supplemental Video 3). This prey item drifted into range, then between frames 1 and 2 made a small appendage motion, which apparently triggered the worm to strike. In this case in the initial strike the proboscis coils around the prey. Several successive strikes over the next few minutes (one is shown in frames 9–12) likely penetrate the cuticle. This large worm takes only a few minutes to completely empty this small prey item (frames 13–15). Notice the whole-body deformation that accompanied proboscis retraction (frame 12). (C) Very similar sequence to (B), corresponds to Supplemental Video 4. Timestamps in ms at first, then in min:sec relative to start of recording. Hollow arrowhead in frame 11 indicates the swallowed eye of the nauplius.

In the simpler capture instances, the nemertean apparently struck prey that merely passed within range, flicking out its proboscis as a chameleon might catch a passing fly. In others, however, there were clear signs that the nemertean used mucus to entrap prey or trigger attacks (Fig. 5B). Many larger individuals of the *Paranemertes* type (e.g. Fig. 2C) trail a visible thread from their posterior end, apparently emanating from the immediate neighborhood of the caudal ciliary cirrus (Fig. 4A and Supplemental Video 2). In smaller individuals mucus threads are not visible even by DIC, but are clearly present from the entrapment of bacteria and debris thereon. This web or cape may extend many body lengths behind the nemertean, and effectively tethers them in glass dishes or on slides. While this tethering may be artifactual, the observation of behavior may not be: when crustaceans (or even forceps or pipettes) blunder into the invisible web, the nemertean often executes a rapid change in behavior, spiraling backward as if to seek the disturber (Fig. 5B and Supplemental Video 6). Once prey is caught in this web, the bundled mucus is often made much more visible by DIC (for example, Fig. 3A).

**Figure 5.**
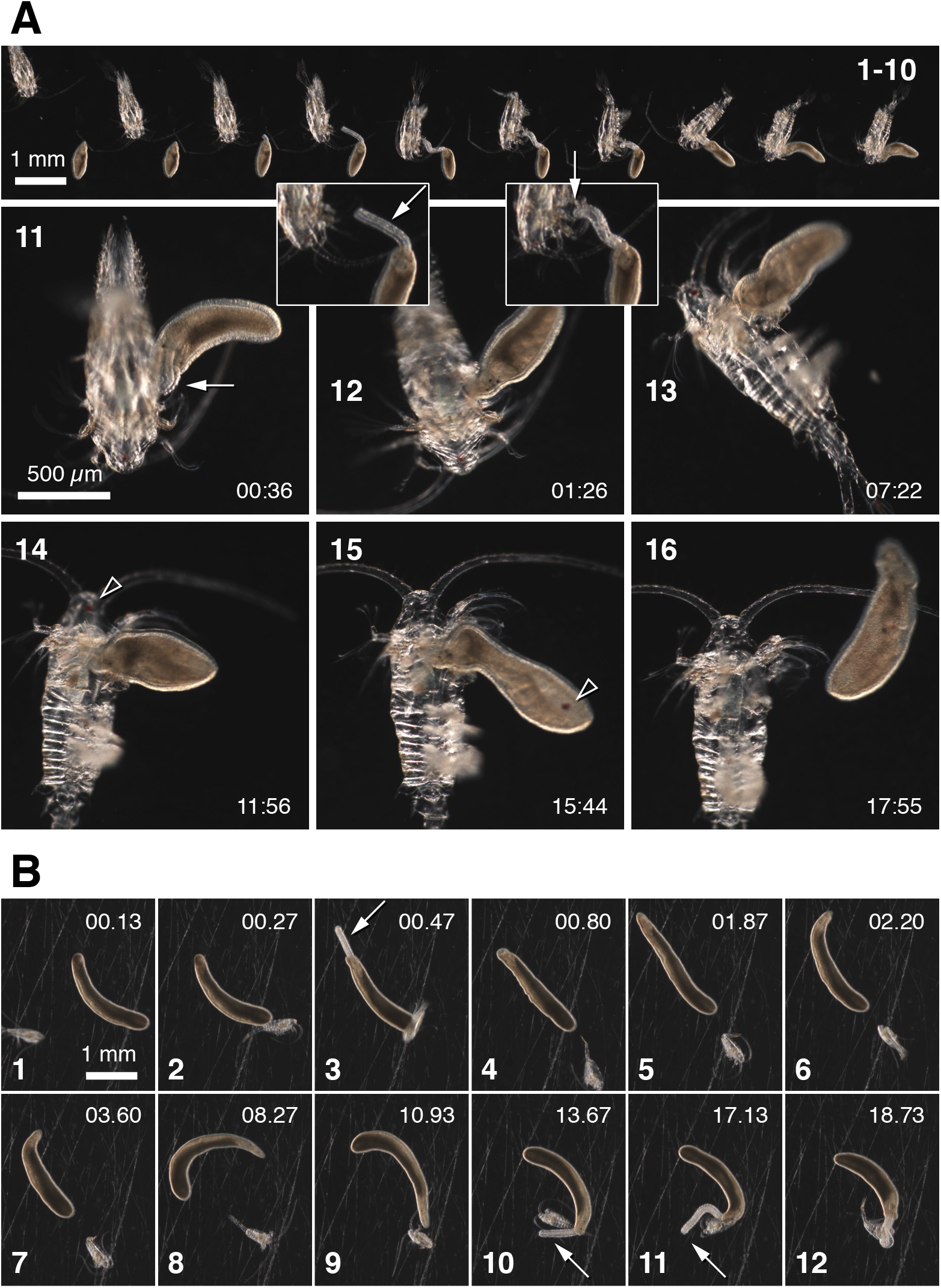
Prey strikes by *Paranemertes californica*. Sequences correspond to Supplemental Videos 5 (A) and 6 (B). The salient features of the sequence summarized in (A) is that, first, the prey, a calanoid copepod, is both highly mobile and several times larger than the predator. Second, as highlighted by insets, the proboscis need not immediately strike for the worm to capture its victim; in this case, the tip of the proboscis (arrow) initially contacted only the head appendages of the copepod, and the stylet may not have penetrated the cuticle until a subsequent strike. In frame 11, the nemertean strikes again; the video shows that the copepod, still alive, reacts violently. Between frames 12 and 13 the pair turned over in the dish. Note the extended thoracopods, which are normally held bundled right next to the body. This posture with thoracopods held out is also achieved by narcotization with tricaine. In frame 13 the nemertean has broken into the cuticle at the site of the strike shown in frame 11, and begun to ingest tissue (note the hollow space next to the nemertean’s head). Over the next ~10 min the nemertean ingested only a portion of the prey (hollow arrowhead in frames 14 and 15 indicates the copepod’s eye, first in its head and then in the nemertean’s intestine). (B) The salient feature of this sequence, which features a very large nemertean that had fed in the lab for many weeks, is that the copepod became entrapped by an invisible web extending posterior to the nemertean. To appreciate this fully requires viewing Supplemental Video 6. Times here are in seconds. The copepod initially darted into view, but was arrested near the posterior end of the nemertean. In frame 3 the nemertean everted its proboscis, but of course the prey was elsewhere. Motion blur and antenna posture in frames 3-7 show that the copepod repeatedly attempted to jump away but was restrained. This well-fed nemertean was apparently in no hurry to feed; over the course of 15 seconds (frames 6-11), it turned around, approached the trapped prey with its anterior end, then struck with its proboscis, missing at least once before finally nailing it.

### 3.3 Identification of suitable prey

Wild-caught larvae conforming to the *Paranemertes* type fed on barnacle and copepod nauplii (Figs. 3, 4B,C), barnacle cyprids, calanoid copepodites and adults (Figs. 1B, 4, 5), decapod zoeas (Fig. 1A), euphausiid nauplii, and cladocerans – essentially any pelagic crustacean we offered, but no non-crustacean prey. Several other wild-caught hoplonemerteans of unknown species were observed feeding on nauplii (the most readily-available offering), but some types never fed in captivity on any offered prey, and eventually dwindled away. One wild-caught hoplonemertean larva of the *Paranemertes* type was observed repeatedly to feed on nereid nectochaetes, but the specimen was unfortunately lost before it could be barcoded to verify its identity. We observed and documented wild-caught larvae identified as *Emplectonema viride* attacking and devouring barnacle nauplii and cyprids (Fig. 6 and Supplemental Video 6). *E. viride* and *Paranemertes*-type larvae are scarce in plankton samples, but are easily recognized and appear reliably in the sub-millimeter size range during late Winter and Spring. Several of these grew rapidly when provided with abundant suitable prey, achieving lengths of several millimeters in lab captivity.

**Figure 6.**
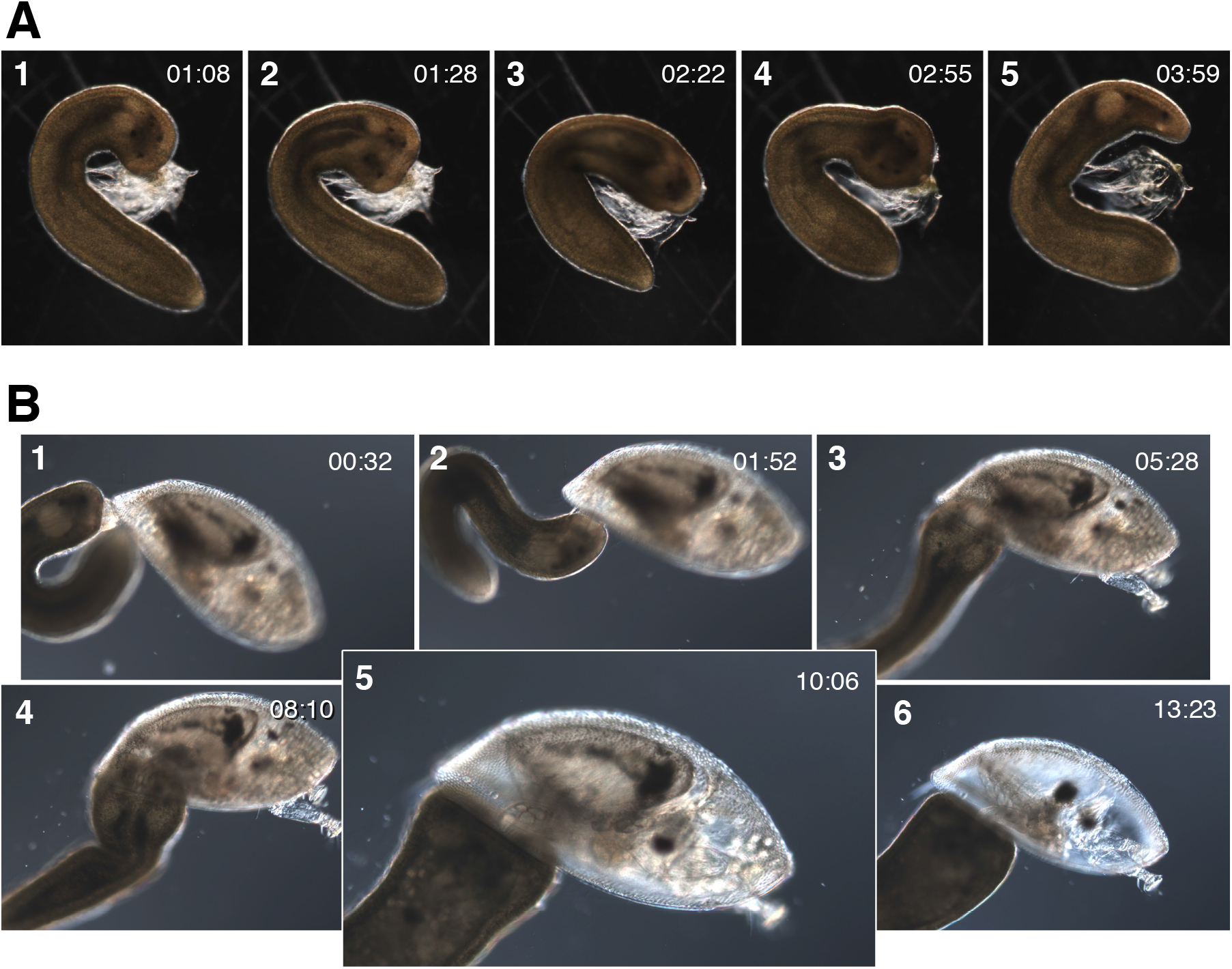
Larvae of *Emplectonema viride* attack and devour barnacle nauplii and cyprids. (A) A relatively large wild-caught larva of *E. viride* eats a nauplius. Times in min:sec from start of recording, which began some minutes after initial attack. In frame 2 the nemertean everted its proboscis again, whereafter it extracted all soft tissues within a few minutes. (B) *E. viride* eating a cyprid (corresponds to Supplemental Video 7). Times in min:sec from start of recording, which commences with the initial attack. In frame 1 the stylet is visible at the center of the short proboscis. In frame 2 the worm stabbed the cyprid again; several successive attacks apparently enabled access through the region behind the thoracopods. At the point where the nemertean had intruded its head under the carapace (frame 3) the cyprid was seemingly still alive, since it moved its antennae actively (although it did not swim). In frames 4-6 the worm drinks in a continuous stream of dissociating tissue; frame 5 is a 1.5x blowup in which lipid granules stream from the cyprid’s head into the nemertean’s mouth.

Repeated trials in which wild-caught *Carcinonemertes epialti* larvae (Fig. 7A,B) were offered numerous potential prey – including decapod, calanoid or harpactacoid copepod, or barnacle eggs, larvae, or even adults – failed to yield any evidence of feeding. In desperation, four *Carcinonemertes* larvae each ~1 mm long were added instead to a small, well-settled plankton subsample. The bowl was then surveyed continuously for nearly two hours by stereomicroscope, before one of the *Carcinonemertes* was spotted with its nose against a small, clear copepod nauplius, which it consumed over the next 10 min. Thereafter a different individual was spotted devouring a copepodite of *Oithona*.

**Figure 7.**
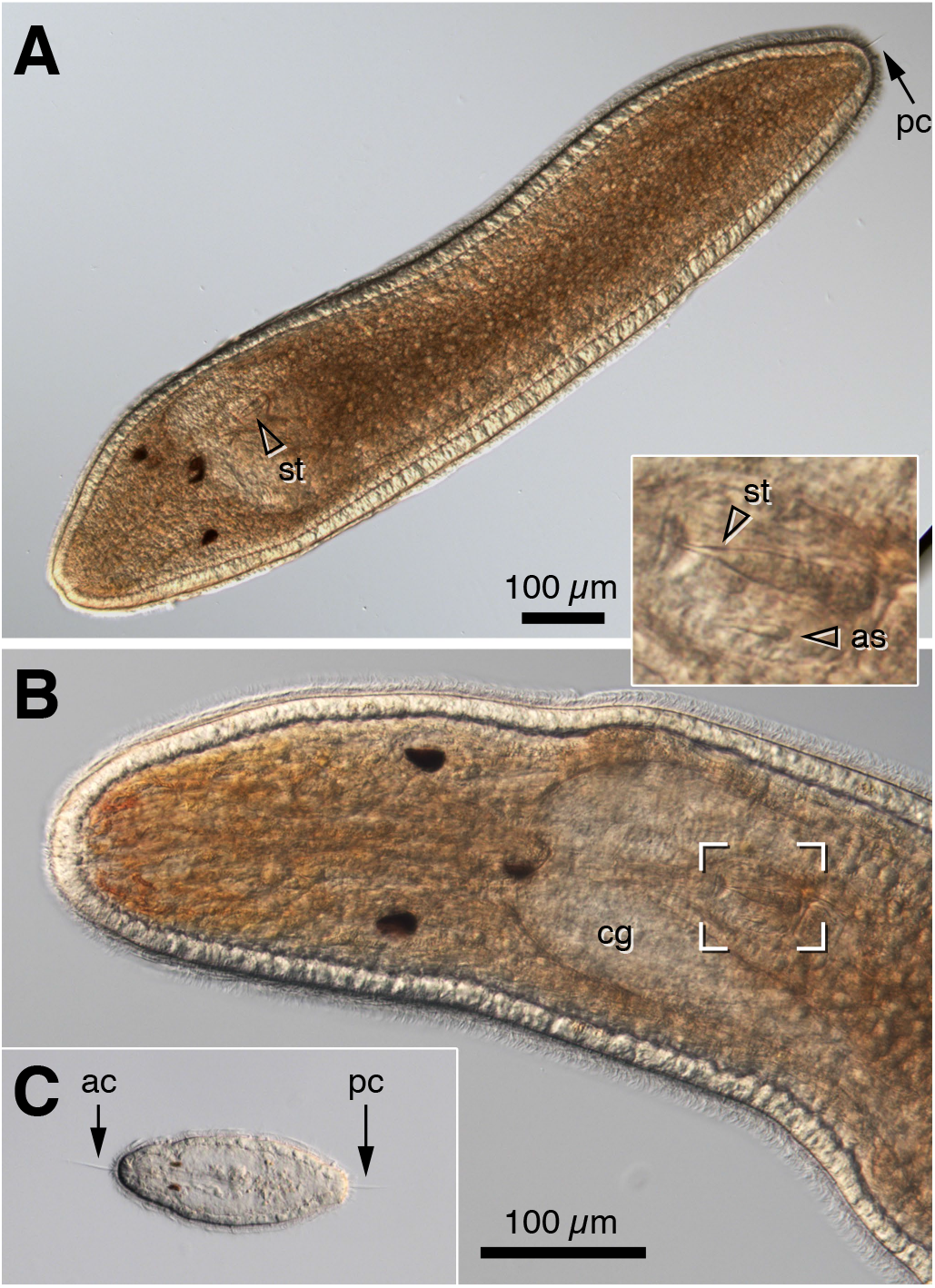
Planktonic larvae of *Carcinonemertes epialti*. (A) A typical wild-caught *C. epialti* larva is ~1 mm long, brick red or orange, has two pairs of submuscular eyes, a single posterior ciliary cirrus (pc, arrow), and a stylet (st, hollow arrowhead) at the end of its short proboscis. (B) Higher-magnification view showing relative arrangement of head parts (one of the medial eyes is out of focus); inset is a contrast-enhanced 2.5x blowup of the indicated region to show stylet and accessory stylet (as); cg=cerebral ganglion. (C) Hatchling *C. epialti* from *Cancer antennarius*. Shown at same scale as (B); the hatchling is roughly the size of the settlement-size larva’s cerebral ganglion. Hatchlings have anterior (apical tuft) and posterior cirrus (arrows; ac and pc), two eyes, no stylet, and are nearly colorless.

### 3.4 Hatchling *Carcinonemertes* grow on a diet of cyclopoid nauplii

Upon this discovery we therefore offered these large planktonic *Carcinonemertes epialti* larvae, which were likely close to competence at the time of capture and had been in captivity for several weeks, an abundance of cyclopoid and other copepod prey, but directly observed only a few more feeding events. However, we had previously obtained hatchling *C. epialti* from the brood of *Metacarcinus magister* (Fig. 7C). These tiny larvae, <80 microns, had been aswim in bowls for weeks and were slowly but surely dwindling in number and size. We hand-selected small *Oithona-type* nauplii from plankton samples and added them to the bowls of *C. epialti* hatchlings, and within minutes observed several latching onto nauplii, then devouring them all or in part (Fig. 8A,B).

**Figure 8.**
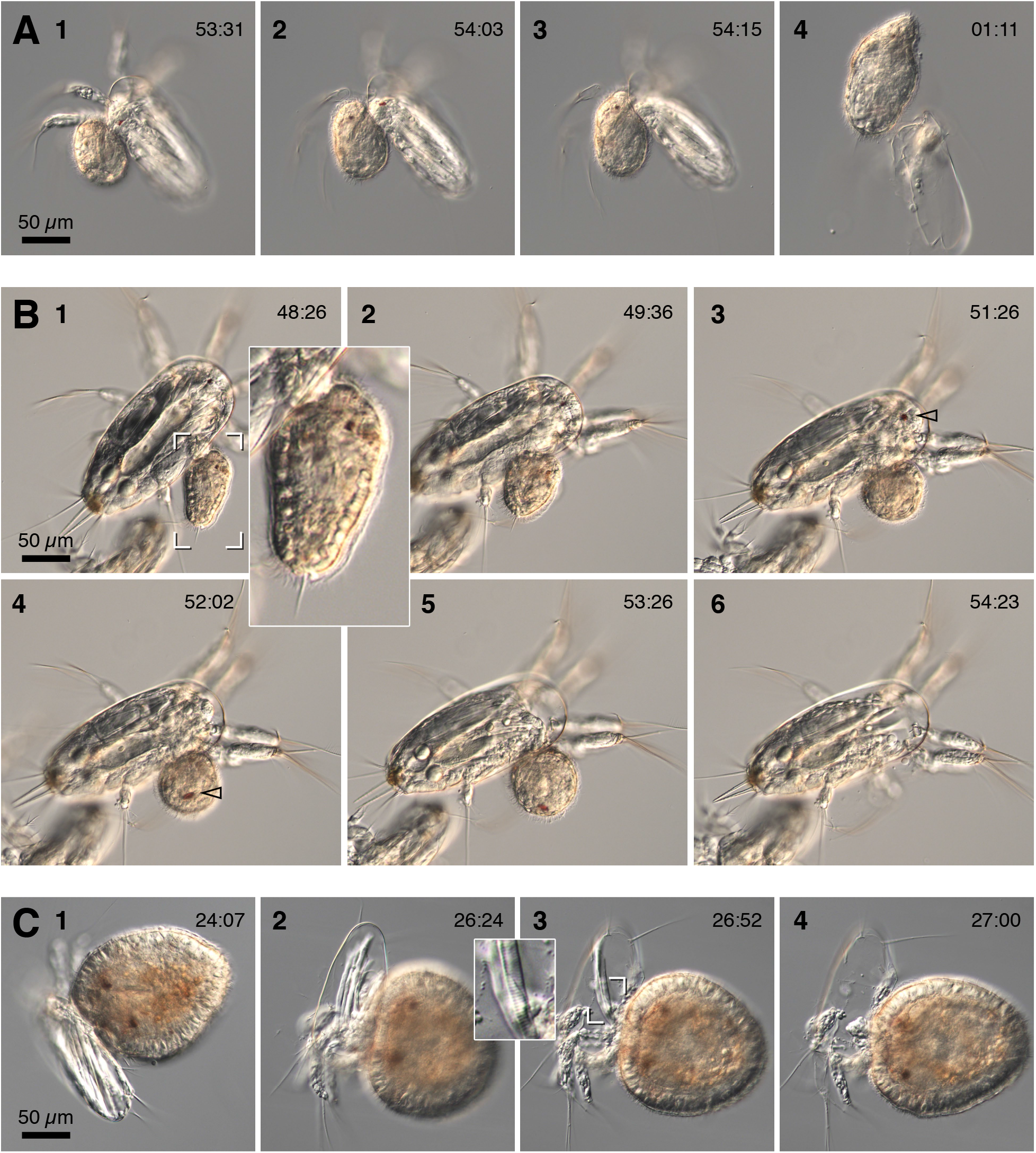
*Carcinonemertes epialti* larvae attack and devour cyclopoid nauplii. (A) Hatchling *C. epialti* from *Metacarcinus magister* encountered wild-caught cyclopoid nauplius in a slide prep and latched on in the armpit region. This represents its first meal. Time stamps are min:sec in clock time. In the first frame show, the worm has already broken through and begun to ingest naupliar tissue; by the second, note that the flesh has been extracted from some naupliar appendages and the naupliar eye is about to be swallowed; by frame 3 the worm has sucked in all the soft tissues of the head, and at this point prey tissue remained seemingly intact. Astonishingly, this particular *C. epialti* hatchling managed to engulf nearly all of the naupliar tissue, expanding greatly in size. (B) A more typical case, also a hatchling from the same batch as (A), evacuated only the head of its much larger victim, leaving a decerebrate but seemingly otherwise intact copepod corpse. From the accumulation of such corpses in bowls with hatchlings, we infer this is a typical outcome. Time stamps are clock time in min:sec. Inset in first frame is a 2.5x blowup of the indicated region, showing the nemertean, in side view, just after latching on to the copepod’s armpit with its mouth. Between frames 3 and 4 the copepod’s eye (hollow arrowhead) was ingested. (C) A lab-reared *C. epialti* larva ~2 months old, consuming a cycploid nauplius in essentially the same way as (A) and (B), albeit somewhat more rapidly. Times are min:sec of clock time. Note the lack of any overt sign that tissue is disrupted before ingestion, in contrast to *Paranemertes californica*. Inset is a 2.5x blowup of the region indicated to highlight that striations remain in a large muscle fiber even as it is swallowed.

In nearly all of dozens of observed feeding instances, the nemertean latched onto the nauplius near the basal joint of the second antenna or maxilla (i.e., the armpit) without an apparent proboscis strike (Fig. 8A,B). Even so, cyclopoid nauplii frequently seemed immobilized or subdued somehow, although they did sometimes thrash or dart away immediately after being attacked by the nemertean; this was occasionally successful, but often the nemertean clung to its victim despite vigorous beating. Notably, although stylets are clearly present in the much larger wild-caught *Carcinonemertes* larvae (Fig. 7A,B), we could not find one in hatchlings or even in animals that had grown considerably in the lab (Fig. 9E). Hence they must either select prey that is passivated for other reasons (approaching molt, perhaps) or they must secrete some sedative.

**Figure 9.**
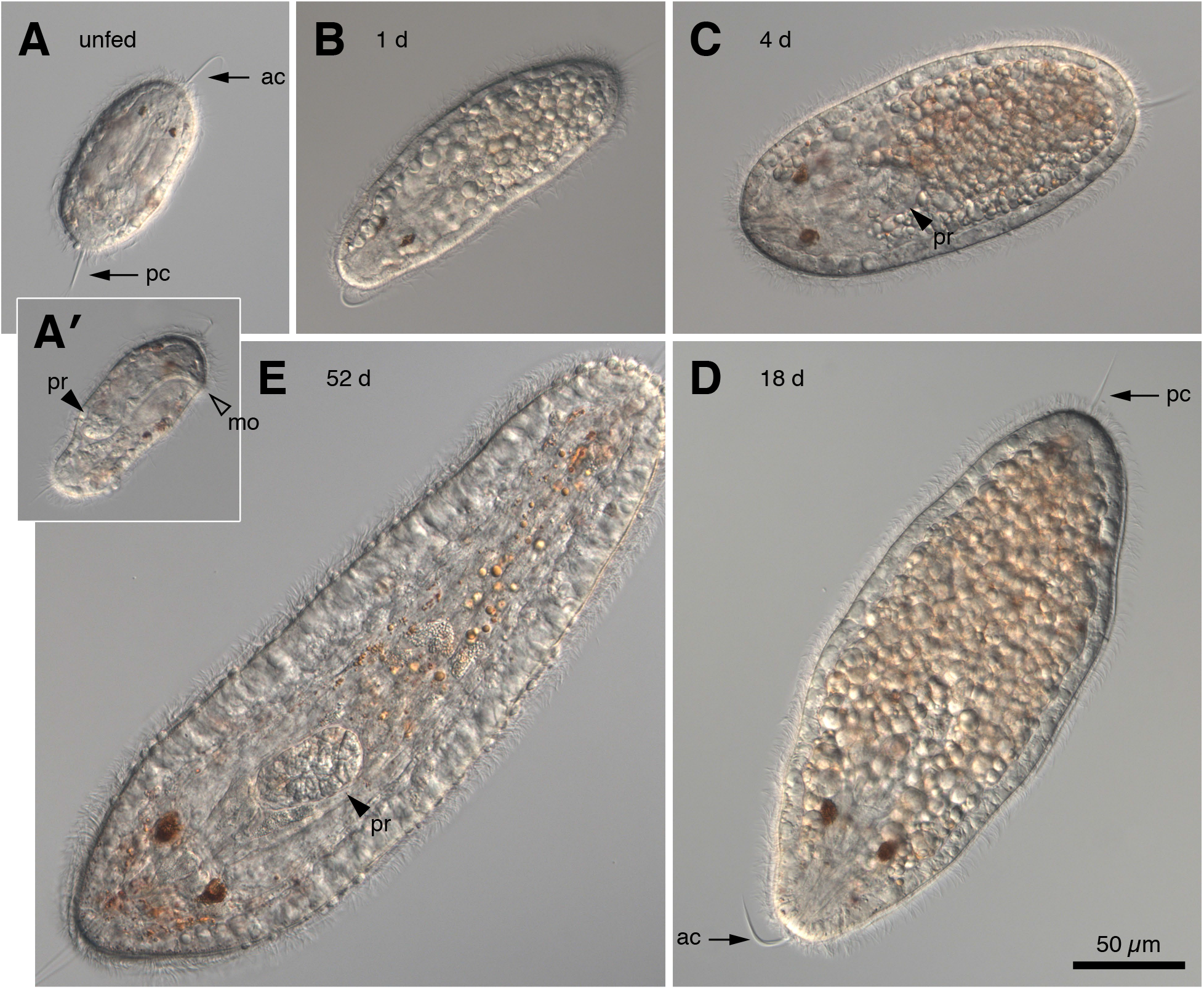
Growth of *Carcinonemertes epialti* in lab culture. All panels shown at the same scale. (A) Unfed hatchling with two eyes, apical tuft and posterior cirrus (arrows, ac and pc), and no distinct granules in the endoderm. (A’) side view of the same individual showing the combined opening of mouth and proboscis (hollow arrowhead, mo) and the location of the posterior end of the as yet unarmed proboscis (arrowhead, pr). (B) A larva from the same batch after eating a single cyclopoid nauplius. Not only is it substantially larger, the endoderm is dominated by globules. (C) After several days with food, larvae have grown substantially and acquired a reddish color to the endoderm; arrowhead shows the position of the end of the proboscis (pr). (D) 18-day-old larva. (E) After nearly two months in culture, starved for the previous week. The endodermal globules have disappeared, and color faded. The proboscis (arrowhead, pr) remains unarmed, and there are only two eyes.

*Carcinonemertes* larvae pumped their own bodies vigorously to break into the cuticle, sometimes dissociating an appendage of their victim. Then they proceeded to suck in tissue, including large, still-birefringent muscle fibers (Fig. 8A,C). In contrast to feeding events involving *Paranemertes-type* or *Emplectonema* larvae, we saw no evidence that tissues were disrupted before ingestion. *Carcinonemertes* hatchlings always targeted prey much larger than themselves (they are ~80 microns; the smallest cyclopoid nauplii we offered were ~150 microns). Even so, some individuals consumed nearly all the soft tissue, expanding greatly to accommodate such an enormous meal (Fig. 8A). More often, however, *Carcinonemertes* hatchlings consumed the head and a few muscles, thereafter abandoning a decerebrate corpse (Fig. 8B). We never saw any evidence of scavenging, and such headless copepods accumulated in culture bowls.

Our attempts to raise *C. epialti* in culture were not, however, entirely successful. Although hatchlings survive for weeks without food, feeding was followed by very high mortality, which we were not able to fully resolve. Some larvae, following feeding, clearly became stuck to glass or to water surface and were damaged irreparably. Coating culture vessels with BSA mitigated some losses, but the larvae grew slowly and continued to die off. Despite substantial growth (Fig. 9) we never saw stylet formation or development of a second pair of eyes, which are found in advanced wild-caught larvae of this species. Examining larvae showed loose surface cells and seemingly related epidermal defects. Whether this is due to the culture conditions, the strain of exceptionally large meals, or the insufficiency of the diet we offered, is unresolved.

### 3.5 *Ototyphlonemertes* larvae also eat copepod nauplii

Amongst the hoplonemerteans that appear in the plankton, larvae of *Ototyphlonemertes* are readily recognized because large individuals have two statocysts in ventral cerebral ganglia (Fig. 10A,B). Like *Carcinonemertes*, these have a very short proboscis. Also like young lab-reared *C. epialti*, they have no stylet at early stages of development. In previous trials we found no evidence of feeding on barnacle nauplii or adult copepods, but inspired by the comparison to *C. epialti*, we offered three captive *Ototyphlonemertes* larvae a choice of copepod nauplii. They promptly captured and fed upon both calanoid and cyclopoid nauplii, entangling these prey, then breaking into the cuticle with repeated eversions of the proboscis, and finally sucking nearly all soft tissues, seemingly intact, out of the cuticle (Fig. 10C).

**Figure 10.**
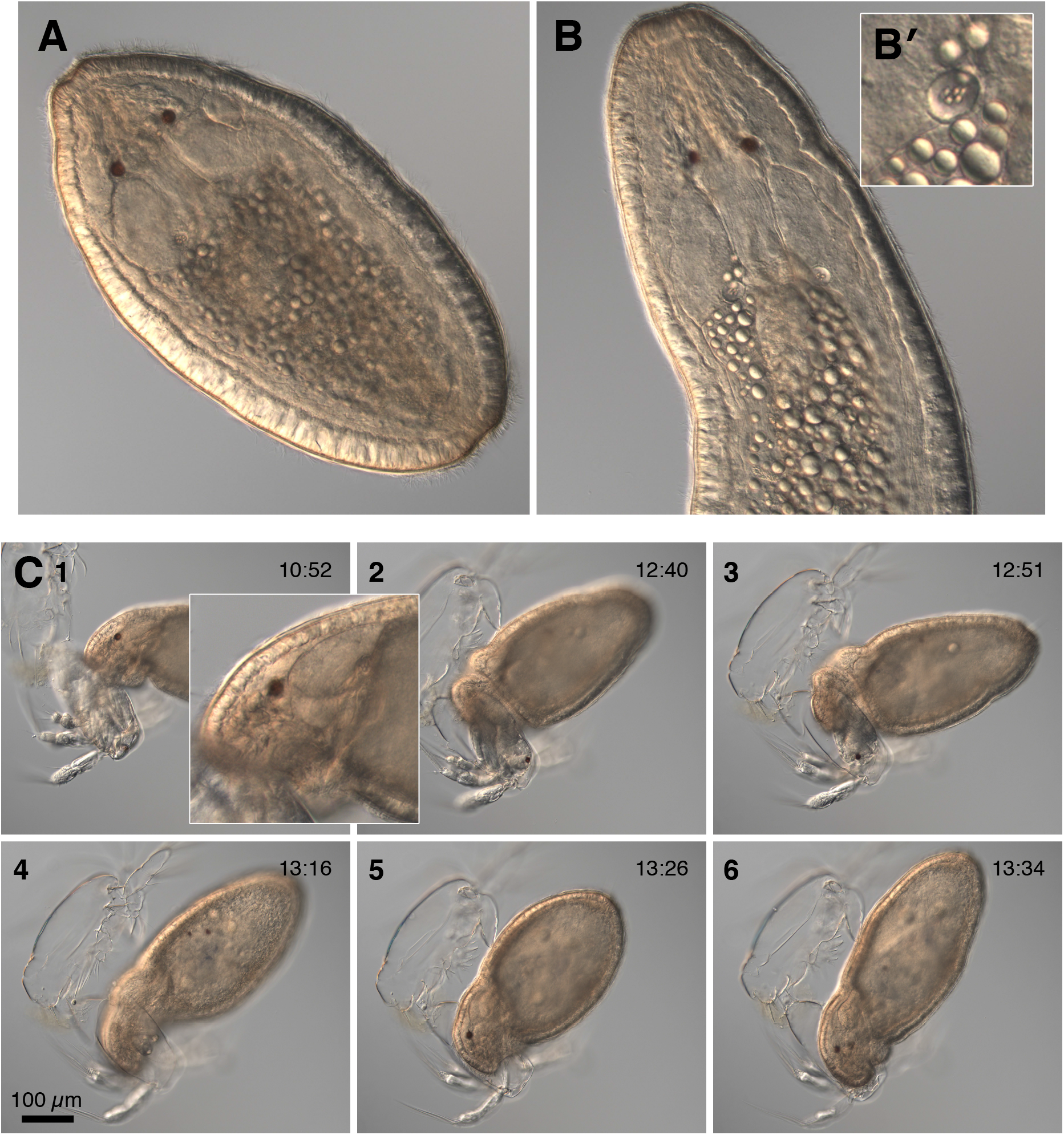
*Ototyphlonemertes* sp. larvae eat copepod nauplii. (A,B) Two wild-caught individuals shortly after collection, both with apparent remains of recent meals in the gut. Inset (B’) is a 3x blowup from a different focal plan to show one of the statocysts located in ventral cerebral ganglia. (C) Image sequence of an *Ototyphlonemertes* larva consuming the second of two copepod nauplii. This worm was noticed after capturing and beginning to eat the first of two entangled nauplii, then transferred to a slide as it began to eat the second. Inset next to frame 1 is a 2.5x blowup of the head from a different focal plane in which a statocyst is in focus (arrowhead). In frame 1 the nemertean had already broken through the dorsal cuticle; by frame 2 the worm had evacuated most of the posterior end of the nauplius; in frame 4, the two red spots in the worm’s gut are the two eyes of its two victims; in frame 5, it sucks the flesh from one of the naupliar antennules.

### 3.6 DNA-based identification of hoplonemertean larvae

Wild-caught hoplonemertean larvae used in this study were identified based on morphology and COI sequences, as belonging to *Emplectonema viride, Carcinonemertes epialti, Ototyphlonemertes* sp., *Paranemertes californica, Paranemertes sp.* (a potentially cryptic sibling species to *P. californica)*, and *Gurjanovella littoralis*, by comparing them to sequences available in GenBank (Table 2).

*Emplectonema viride* STIMPSON, 1857, a barnacle predator, is one of the most common intertidal hoplonemertean species found along the West Coast of North America from Alaska to California. Larvae of *E. viride* can be easily identified by their green color (Hiebert, 2016). COI sequences from these larvae were 100% identical to GenBank sequences of *Emplectonema* sp. 1 derived by the members of the Maslakova lab from morphologically identified adults collected in Charleston, OR, and now known to belong to *E. viride* (Supplemental Figure 1). *E. viride* was briefly described by Stimpson (1857) from San Francisco Bay, CA, and later redescribed by Griffin from Alaska and Puget Sound, WA (1898). This species was synonymized with the Atlantic look-alike *Emplectonema gracile* by Coe (1901) and has been referred to as *E. gracile* for over a hundred years (e.g. Roe et al., 2007). Hiebert (2016) pointed out that the West coast species is distinct from *E. gracile*, and submitted sequences to GenBank under the temporary name *Emplectonema* sp. 1. The species is currently being redescribed as *Emplectonema viride* Stimpson, 1857 (Mendes et al., in press).

*Carcinonemertes epialti* COE, 1902 is a parasite and egg predator of decapod crustaceans, which is known to occur in its adult form along the Pacific Coast of North America on crabs of various species. The infected species include the cancrids *Cancer antennarius, Cancer anthonyi, Cancer jordani, Cancer oregonensis, Cancer productus*, the grapsids *Hemigrapsus nudus, Hemigrapsus oregonensis* and *Pachygrapsus crassipes*, the majids *Pugettia producta*, and the portunids *Euphylax dovii* and *Randallia ornata* (Gibson, 1995), as well as the introduced European green crab *Carcinus maenas* (Torchin et al., 1996). A species described as *Carcinonemertes errans* Wickham, 1978 is reported to occur only on *Metacarcinus magister* (the Dungeness crab). *Carcinonemertes errans* differs from *C. epialti* in that the adults move freely among the crab’s egg mass, and do not secrete a protective sheath (Wickham, 1980). However, COI sequences of *Carcinonemertes* collected by us from *Metacarcinus magister, Pugettia producta*, and *C. maenas* are <1% divergent from each other, or from sequences of *Carcinonemertes* larvae collected by us from plankton in Charleston, OR during this study (Table 2) and in previous years (see Supplemental Figure 1). This suggests that *C. errans* Wickham, 1978 and *C. epialti* Coe, 1902 represent the same species, and that the difference in sheath-building behaviour may be related to the host biology. *Carcinonemertes epialti* has priority over *C. errans*, according to the rules of the International Code of Zoological Nomenclature (Article 23, ICZN 1999). Hence, all specimens of *Carcinonemertes* used in this study, including those collected from *M. magister*, are identified as belonging to *C. epialti*.

Advanced larvae of the interstitial genus *Ototyphlonemertes* can be easily identified morphologically by a pair of statocysts in the ventral cerebral ganglia (as in adults), as well as a pair of ocelli, which the adults lack (Chernyshev, 2000; Hiebert, 2016). COI sequences derived from such larvae were nearly identical to each other and most similar to the sequences of *Ototyphlonemertes santacruzensis* and *Prosorhochmus claparedii* available in GenBank, though none were a species-level match, ranging from 85% to 82% in sequence similarity. On the ML tree our *Ototyphlonemertes* sequences did not group with either species, but formed a separate clade (Supplemental Figure 1).

*Paranemertes californica* COE, 1904 is known from intertidal soft sediments (sand) from Monterey Bay, CA to Ensenada, Mexico (Coe, 1940; Roe et al., 2007), and we have collected a single adult of this species at Sunset Bay near Charleston, OR in 2012 (Maslakova, unpublished). COI and 16S sequences derived from this individual (KU197614, KU197273) served as the basis for identification of previously published larval sequences (Hiebert, 2016) and the larval sequences from this study. The main caveat is that GenBank COI sequences identified as *P. californica* (Hiebert, 2016) form two distinct subclades, with sequence divergences (uncorrected p-distance) within clades of <1%, and interclade divergences of 5.5-6.4%, which may indicate the presence of two cryptic species in a species complex, as these divergences are very close to the upper limit of the observed barcoding gap (4-5%) for COI sequences in Nemertea (Sundberg et al., 2016). While samples of *P. californica-like* larvae from 2019 and 2020 (21III19-2, 21III19-5, 21III19-8, 21III19-9, COIMB2047-Miriam, COIMB2048-Anton, COIMB2049-Victor and COIMB2117-Hester) belong to the subclade that includes the morphologically identified adult of *P. californica* (KU197614), samples N17, N97, N60 and COIMB2046-Elvira, collected in 2013 and 2020 belong to the other subclade (Table 2, Supplemental figure 1), and thus may represent a cryptic sibling species, hence we refer to them as *Paranemertes* sp.

Finally, one of the specimens (COIMB2116-Alfonse) with similar morphology to larvae from the *Paranemertes californica* species complex mentioned above was later identified as *Gurjanovella littoralis*. The COI sequence from this specimen is 99% match to a larva of *G. littoralis* (KU197600) also from Oregon (Table 2, Supplemental figure 1) and 95% match to the sequence of adult *G. littoralis* (AJ436904) collected by SAM from the White Sea, Russia (where this species was originally described). Although there are no published records of *G. littoralis* occurring in the NE Pacific, SAM has previously collected an adult of this species at False Bay, San Juan Island, WA (Maslakova, unpublished obs.). Notably, this is the only successfully-barcoded larva that consistently trailed a prominent “fishing line” thread of mucus from the posterior end (depicted clearly in Fig. 4A); future sampling may reveal whether this is a species-specific trait or, as we originally thought, a developmental stage.

### 3.7 DNA-based identification of stomach contents of wild-caught hoplonemertean larvae

DNA-barcoding of hoplonemertean larval samples using a combination of a 16S rRNA universal forward (16S-ARL) and Eucarid-specific (16S-CruR) primers (Table 1), allowed us to amplify crustacean sequences from four wild-caught hoplonemertean larvae identified here as belonging to the *Paranemertes californica* species complex (Table 2, Supplemental Figures 1 and 2). A sequence identical to that of a pinnotherid crab *Scleroplax littoralis* (EU934975) was isolated from *P. californica* larva 21III19-5 collected in 2019. Sequences identical to that derived from a pinnotherid crab *Scleroplax faba* (EU934976) were isolated from two different larvae (N17 and N97) collected in 2013 and identified as *Paranemertes* sp. within the *P. californica* species complex. The two pinnotherid species formerly known as *Pinnixa faba* (Dana, 1851) and *Pinnixa littoralis* Holmes, 1894 were recently placed within the genus *Scleroplax* by Palacios Theil and Felder (2020). Finally, a sequence likely belonging to a cancrid crab was isolated from another larva (N60) collected in 2013 and identified as *Paranemertes* sp. from *P. californica* species complex. The crustacean sequence in question was 92.7% identical to *Cancer pagurus* (FM207653), and nested within a clade of similar sequences belonging to *Cancer* spp. These prey sequences represent instances of hoplonemertean larval predation in the wild, as opposed to those observed in the laboratory.

### 3.8 DNA-based identification of prey remains from laboratory feeding

In addition to sequences of prey isolated from larvae that fed in the wild, we were able to obtain sequences from remains of some of the prey items hoplonemertean larvae fed upon in the lab (Table 2, Supplemental Figures 1 and 2). In one instance we isolated a COI sequence 99.5% identical to that of *Centropages abdominalis* (HQ966505) from the remains of a copepod fed upon by the larva of *Paranemertes californica* (21III19-2) using universal COI primers (Table 1). In another instance, we obtained a 16S rRNA sequence identical to that of the European green crab, *Carcinus maenas* (FM208763 and others), from the remains of a wild-caught zoea larva fed upon by the larva of *Paranemertes californica* (21III19-8) using a combination of a 16S rRNA universal forward (16S-ARL) and Eucarid-specific (16S-CruR) primers (Table 1). *Carcinus maenas* has been introduced to the Pacific Coast of North America in the late 1980s, and is apparently facing some pressure from the local nemertean predators and parasites (Torchin et al., 1996). Furthermore, we identified remains of two barnacle nauplii fed upon by the lab-reared larvae of *Emplectonema gracile* as *Balanus glandula* (99.7% match to EF552057 and others, and those of numerous copepod nauplii fed upon by the larvae of *Carcinonemertes epialti* as *Oithona* sp. Sequences from these copepod nauplii did not have a species-level match to any sequences in GenBank, the closest match being *Oithona similis* (around 84% similarity to JN230862 and others).

## 4. Discussion

We report evidence that at least six species of hoplonemertean larvae practice macrophagous carnivory. From this finding the clear inference is that these larvae make a living in the plankton by taking prey similar to those we observed them to feed on in the lab. The question naturally arises, however, are these really planktonic larvae? Both words in this term are questionable. Certainly they swim and are caught in plankton tows, but they could appear in the plankton spuriously; indeed when captive in bowls they spend a large fraction of their time either sedentary or swimming in close association with surfaces. But the clearest reason to think that they are really planktonic is the nature of the prey they eat.

The second word in question is whether they qualify as larvae, as opposed to juveniles. These nemertean young possess few traits that distinguish them from juveniles, and lack few that characterize adults. In particular, they possess a proboscis armed with stylets (except for young *Ototyphlonemertes* and *Carcinonemertes*) and cerebral organs (although some species apparently lack them at early stages). Subtle changes constitute a sort of metamorphosis: rearrangement or loss of eyes, cirri, epidermal cells, and changes in ciliary length or behavior (e.g. Maslakova and von Döhren, 2009; Maslakova and Hiebert, 2014; Dunn and Young, 2014; Mendes et al., in prep.). The subtlety of these changes argues that from an embryological point of view, these are direct developers: they are planktonic juveniles whose prey capture and preferences are similar to the adults they will eventually become. However, ecologically they are larvae: they occupy a different habitat and exploit a different resource therein, compared to the adults they will become, and – assuming they really live in the plankton – they experience the same opportunities for dispersal or site selection that any long-lived planktotrophic larva does.

It is remarkable that any soft-bodied ciliated larva should successfully prey upon pelagic crustaceans, especially copepods, which are relatively well-defended, are among the fastest of swimmers, and are acutely sensitive to water motion (Kiørboe et al., 1999; Yen, 2000; Buskey et al., 2002). Furthermore, concentrated plankton samples tend to convey a misleading impression of the potential frequency of encounters amongst small pelagic organisms. While copepods and barnacle nauplii are among the most abundant zooplankters, plankton is usually dilute. Our video recordings have not given us any insight into whether hoplonemerteans might detect and swim toward prey at a distance. Indeed, in most recordings, the predator either blunders into prey or strikes at it within the short radius set by the proboscis. Therefore, it is of interest to consider whether and how a comparatively slow swimmer can hope to encounter enough to eat.

Hoplonemertean larvae can swim on the order of 0.1-1 cm/sec., and tend to do so in a steady helical motion or in widening spirals. For small individuals the apparent encounter radius of a few hundred microns suggests that these animals sample a water volume on the order of a microliter per second, or a few liters per hour (if their path is not retraced). Copepods and their nauplii routinely exceed a density of 1 per liter in surface waters (e.g., Galienne and Robins, 2001; Turner, 2004). Not every encounter or strike is successful, but one success may gain the nemertean considerably more than its own body size. Perhaps, like pythons, their meals are so large that they need be only infrequent. Hence it seems likely that with even modest capture efficiency, a relatively slow-swimming ciliated blob might make a good living.

Another possibility is that instead of searching for prey by swimming, hoplonemertean larvae may self-tether using mucus either to associate themselves with debris or suspended particles or to sail in the water column. Like web-building spiders, they may entrap prey in mucus threads, thus expanding their reach and slowing down fast-moving prey items enough to approach and evaluate them. Even without relying on mucus threads for entrapment, a sit-and-wait approach by a sailing predator equipped with a fast strike may be just as effective as a swimming search, since the high motility of crustacean prey ensures that each potential prey item is likely to visit a large volume per unit time.

For the species we studied we think we can tentatively exclude one other possible strategy: that these nemertean larvae are actually infaunal. We initially considered this possibility for *Carcinonemertes* because of our long inability to identify acceptable food for them. But none exhibit any inclination to bury themselves in sand or sediment when offered the opportunity to do so, and all swam actively at least some of the time, and furthermore we never observed feeding on harpactacoid copepods or other benthic potential prey. An interesting possibility, however, is raised by the chance discovery of *Emplectonema viride* juveniles within the mantle cavity of lepadomorph barnacles (Eric Sharman, George von Dassow, and Svetlana Malsakova, unpublished observations.). It is not yet known what they are doing there or how consistently they are present, but it makes us curious whether some small planktotrophs take advantage of large-bodied filter feeders to concentrate plankton for them. At least one hoplonemertean genus (*Malacobdella*) lives this way as adults. So far, however, casual surveys inside possible hosts (clams, mussels, ascidians) have not turned up any clandestine population of hoplonemertean larvae lurking within.

Finally, our discovery has far-reaching implications for animal life history studies and the evolution of developmental mode. Classically, zoologists stereotype animal larvae into planktotrophs that feed and grow on suspended primary producers (unicellular phytoflagellates, primarily) versus lecithotrophs that depend on maternal provisions to build a viable juvenile directly from the resources in the egg (Thorson, 1950; Jägersten, 1972; Strathmann, 1985; Fell, 1997). The nemertean pilidium larva is a canonical example of the former, whereas the ascidian tadpole epitomizes the latter. These examples underscore that either larval type, planktotroph or lecithotroph, might differ morphologically from the juvenile and transit a potential dispersal phase before undergoing metamorphosis and settlement. Indirect-developing planktotrophic larvae, like the pilidium, typically deploy larval-specific food-collecting devices which are lost or consumed at metamorphosis (Jägersten, 1972; Strathmann, 1985; specific larval types, pilidium: Maslakova, 2010a; actinotroch: Temereva and Malakhov, 2015; mitraria: Wilson, 1932; echinoderms: Müller, 1850; Cameron and Hinegardner, 1978). Lecithotrophic larvae too, like the ascidian tadpole, may have larval-specific apparatus for dispersal and site selection (ascidian: Cloney, 1982; bryozoan coronate: Reed, 1985). Planktotrophs that feed on phytoflagellates may be able to afford to have many small eggs, whereas lecithotrophic equivalents may need to make much larger eggs to be able to form a viable juvenile (Vance, 1973; Strathmann, 1985; Emlet et al., 1987), except perhaps for animals like ascidians and bryozoans, whose juveniles themselves live on nanoplankton. In contrast to both of these explicitly larval stereotypes, when zoologists find that an animal’s egg hatches without special larval traits, it is natural to infer that the hatchling is a juvenile about to live more or less like the adult. Such has been the supposition for hoplonemerteans (e.g. Norenburg and Stricker, 2002), for polyclads (Shinn, 1987; Martín-Durán and Egger, 2012), and for various annelids (Wilson, 1991) and other invertebrates. Our discovery that at least some hoplonemertean larvae live in the plankton as macrophagous carnivores illustrates an additional strategy. A companion paper on pelagic polyclad larvae makes a similar case (von Dassow and Mendes, submitted). Palaeonemertean larvae live similarly (e.g. Maslakova and Hiebert, 2014). Other instances across the invertebrate world, including some anthozoan planulae and annelid trochophores, may not be so exceptional.

## Acknowledgements

CM was supported by São Paulo Research Foundation (FAPESP) grant 2019/10375-8.

---

**Supplemental figure 1.**
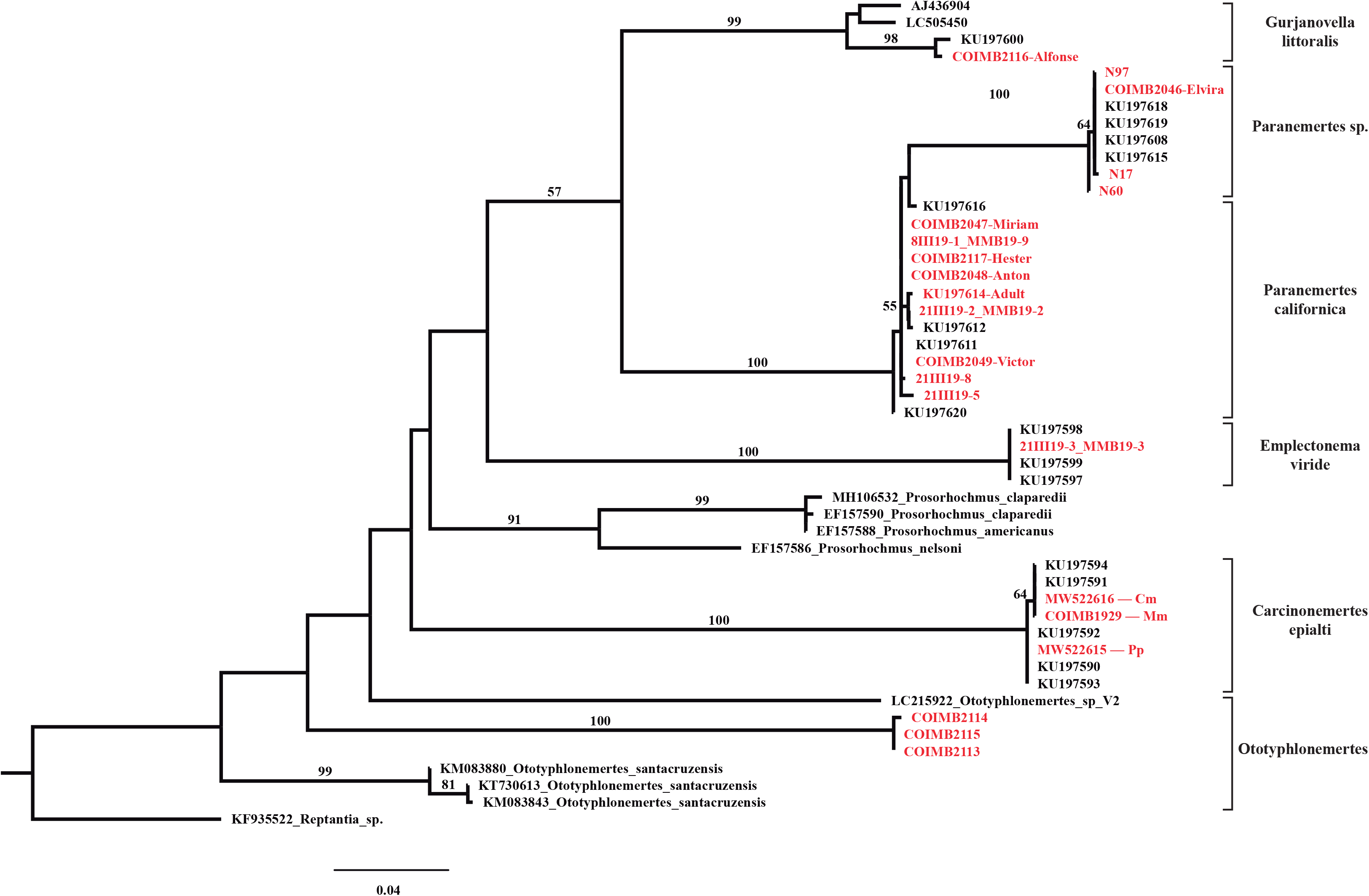
A Maximum Likelihood phylogeny constructed using RAxML from COI sequences of hoplonemertean larval samples and three of their respective closest matches from GenBank (InL = −3631.08). Bootstrap support values below 50% are not shown. Sequences shown in red are obtained in this study, in black are comparison sequences from GenBank. For *Carcinonemertes epialti*, crab species from which specimens were collect is shown as *Mm=Metacarcinus magister, Cm=Carcinus maenas, Pp=Pugettia producta*.

**Supplemental figure 2.**
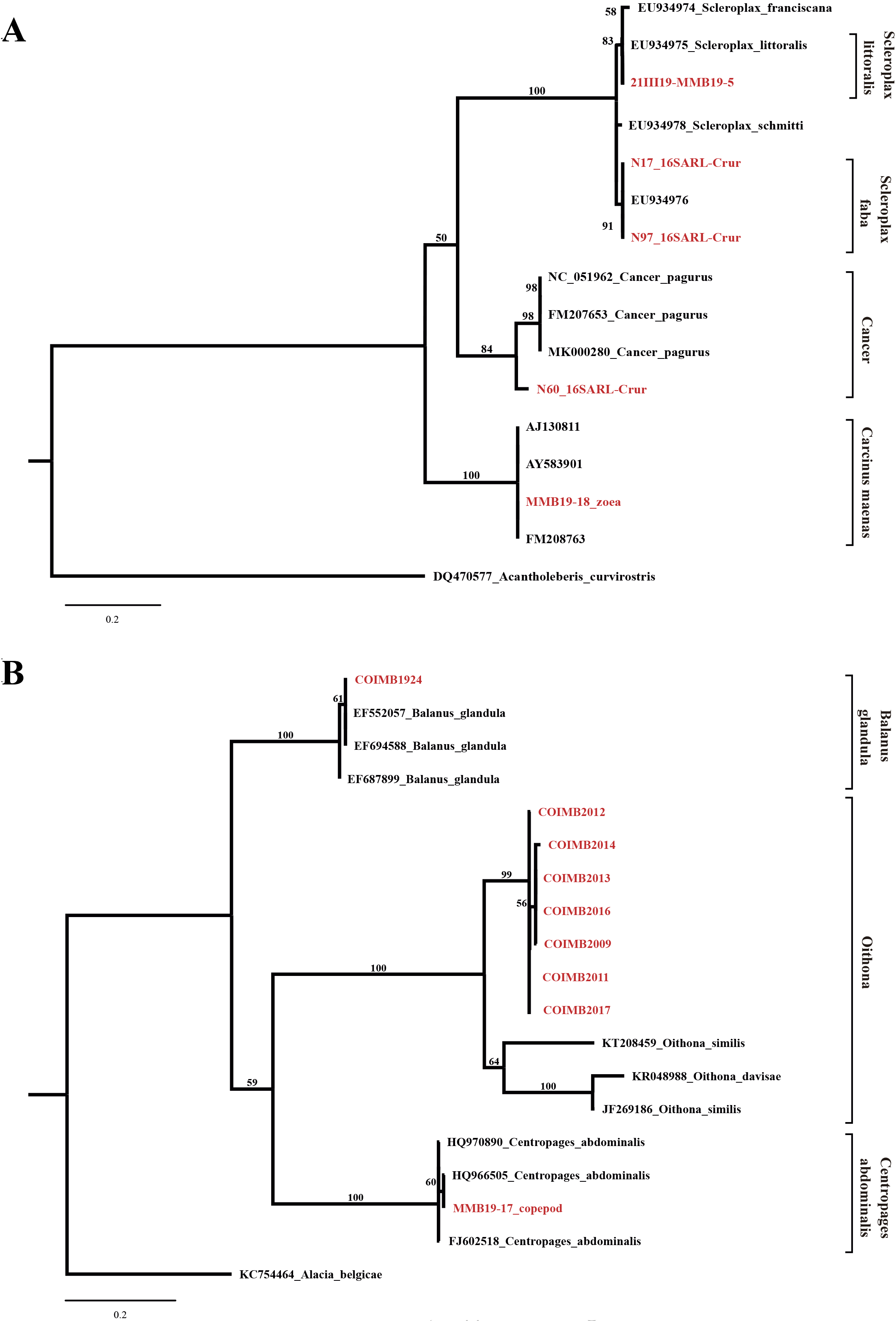
Maximum Likelihood phylogenies (A: 16SrRNA, lnL = −1604.2619; B: COI, lnL = −3743.2685) constructed using RAxML from prey sequences amplified from hoplonemertean larval samples (i.e. gut contents) or from remains of prey items consumed by hoplonemertean larvae in the lab (red). Comparison sequences from GenBank (three closest matches) are in black. Bootstrap support values below 50% not shown.

